# Metabolism of epigenetic ribonucleosides leads to nucleolar stress and cytotoxicity

**DOI:** 10.1101/2025.06.11.659152

**Authors:** Xuemeng Sun, Anita Donlic, Jacob A. Boyer, Neal Reddy, Clifford P. Brangwynne, Joshua D. Rabinowitz, Ralph E. Kleiner

## Abstract

Post-transcriptional RNA modifications are ubiquitous in biology, but the fate of epigenetic ribonucleotides after RNA turnover and the consequences of their metabolism and misincorporation into nucleic acids are largely unknown. Here we explore the metabolism of epigenetic ribonucleosides in human cells by studying effects on cell growth, quantifying misincorporation into cellular RNAs and identifying metabolic regulators, and exploring phenotypes associated with cytotoxicity. We find that bulky N^6^-modified adenosines (i.e. i^6^A) exhibit high levels of cytotoxicity and RNA misincorporation, whereas cells dramatically restrict the misincorporation of small N^6^-modified adenosines (i.e. m^6^A), partly through sanitization by enzymatic deamination. Epigenetic ribopyrimidines also exhibit cytotoxicity, mediated primarily by nucleoside kinase UCK2, but only at much higher concentrations than ribopurines. We further characterize the effects of cytotoxic ribonucleoside metabolism on nucleolar morphology and protein translation. Taken together, our work provides new insights into the metabolism of epigenetic ribonucleosides and mechanisms underlying their cytotoxicity to cells.

## Introduction

Nucleotides are essential for life, serving as the building blocks for RNA and DNA. Beyond their role in nucleic acid synthesis, nucleotides also participate in diverse signaling pathways. It is well established that imbalances in the nucleotide pool can disrupt cellular processes and contribute to various human diseases^1–5^.

Cells generate nucleotides through two primary biosynthetic routes: *de novo* synthesis and salvage pathways. The *de novo* pathway generates nucleotides from basic precursors including amino acids, sugars, and bicarbonate. Conversely, salvage pathways recycle nucleobases or nucleosides derived from RNA/DNA turnover or from external sources, such as dietary nucleic acids and cellular degradation products^2,6^. These two pathways are interconnected, allowing cells to compensate for inhibition of one pathway by upregulating the other^3,7^. Notably, the relative contributions of *de novo* synthesis and salvage vary significantly across tissues^7^.

Because the salvage pathway recycles nucleotides derived from nucleic acid turnover, this pathway is exposed not only to canonical nucleotide structures, but also modified epigenetic nucleotides found in RNA/DNA. In particular, over 150 distinct chemical modifications are known on cellular RNA, spanning ribosomal RNA (rRNA), transfer RNA (tRNA), messenger RNA (mRNA) and other noncoding RNAs^8,9^. These modifications are highly conserved across all kingdoms of life and are involved in regulating fundamental biological processes through modulation of RNA structure, base pairing, and protein-RNA interactions^8,10,11^. Dysregulation of RNA modifications has also been implicated in numerous human diseases, including cancer and neurological disorders^8,12,13^.

Among the diverse collection of RNA modifications, some are restricted in their abundance or exotically structured and likely incompatible with metabolism^8,12^, and therefore unlikely to make substantial contributions to cellular nucleotide pools. In contrast, a number of modified RNA nucleotides are broadly conserved and abundant (present in concentrations ∼1-10% of canonical nucleotides) and structurally similar to canonical nucleotides. These include pseudouridine (Ψ)^8^, dihydrouridine (D)^8^, 5-methyluridine (m^5^U)^8^, 5-methylcytidine (m^5^C)^8^, and N^6^-methyladenosine (m^6^A)^11^, among others. Despite the prevalence of RNA modifications, the metabolic fate of modified RNA nucleotides after RNA turnover remains largely unknown. Whether specialized pathways exist to distinguish and process these structures, and what cellular consequences arise from their improper metabolism, are open questions (Figure 1A).

**Figure 1.**
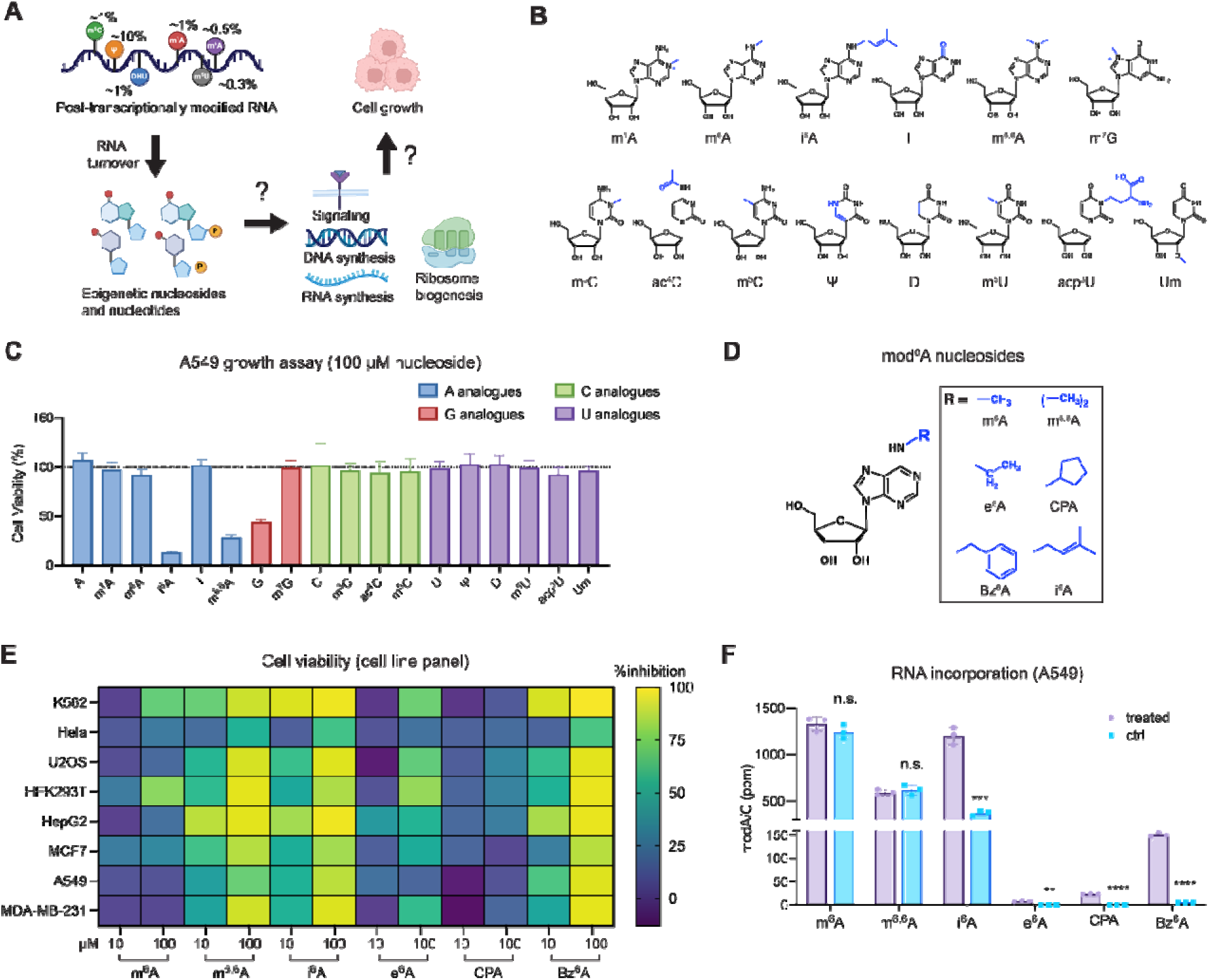
Metabolism of epigenetic ribonucleosides. **(A)** Scheme of potential fates of epigeneti ribonucleosides that originate from post-transcriptionally modified RNA. **(B)** Structures of modified epigenetic purines and pyrimidines used in this work. **(C)** Cytotoxicity of epigenetic ribonucleosides (100 µM for 72 h) in WT A549 cells. Cell viability was measured by an MTS-based assay and plot represent mean normalized cell viability ± s. d. (*n=9*; three independent biological replicates with three technical replicates for each). **(D)** Structures of modified N^6^-adenosine analogs used in this work. **(E)** Screening of N^6^-modified adenosine cytotoxicity in a panel of human cell lines. Heatmap shows percentage growth inhibition with 10 µM or 100 µM nucleoside treatment for 72 h, corresponding to Supplementary Figure 1C-J. Cell viability was measured by an MTS-based assay and represents mean normalized by untreated cells. (*n*=9 with three technical replicates for each of the three independent biological replicates). **(F)** RNA modification levels in total RNA after treatment with N^6^-modified adenosine analogs. WT A549 cells were treated with 10 µM nucleoside for 24 h. Modified nucleotide levels were quantified by nucleoside LC- QQQ-MS. Data are mean ± s.d. (*n*=3). Multiple unpaired t-test were performed between control and treated. Adjusted p values for e^6^A: p=0.0024; for CPA: p=0.000001; for: i^6^A: p=0.00052; for Bz^6^A: p=0.000002.

Previous studies investigating the metabolism of epigenetic nucleosides have focused primarily on deoxyribonucleosides. Salvage and misincorporation of the major eukaryotic epigenetic DNA modifications, 5-methylcytosine and its oxidized derivatives, is controlled by multiple metabolic enzymes in the pyrimidine salvage pathway. Direct salvage of 5-methyl-2’-deoxycytidine, the most abundant mark, is restricted by cytidine/uridine monophosphate kinase (CMPK1) and cytidine deaminase (CDA)^14^. CMPK1 also restricts salvage of 5-hydroxymethyl-2’-deoxycytidine (5hmdC) and higher oxidation states (i.e. formyl, carboxy)^14^, however deamination to 5- hydroxymethyl-2’-deoxyuridine (5hmdU) in cells with high CDA activity results in DNA misincorporation, DNA damage, and cytotoxicity. Cells additionally express 2’-deoxynucleoside 5’-monophosphate N-glycosidase (DNPH1), a sanitization enzyme that processes 5hmdU monophosphate (5hmdUMP) and prevents its DNA misincorporation^15^.

An analogous sanitization pathway has been proposed for the abundant epigenetic ribonucleoside N^6^-methyladenosine (m^6^A). Studies in *Arabidopsis thaliana* and human cells^16^ identified N^6^-methyl-AMP deaminase (ADAL) as an enzyme that deaminates N^6^-methyl-AMP (m^6^AMP) and N^6^-methyl-2’-deoxyAMP (m^6^dAMP)^17^ into their corresponding inosine derivatives. These findings suggest that ADAL prevents metabolism and misincorporation of N^6^-methyladenosine into DNA^17,18^ and RNA, however direct evidence for a role in preventing RNA misincorporation is lacking^16^. Additional studies of epigenetic ribonucleosides have found that extracellular N^6^-isopentenyladenosine (i^6^A) is cytotoxic to mammalian cells and can be incorporated into cellular RNA^19^; in *Arabidopsis*, loss of nucleoside hydrolase 1 (NSH1) correlates with accumulation of 5-methyluridine (m^5^U) into mRNA and reduced seedling growth^20^, however mammalian NSH1 homologs have yet to be identified. Overall, the metabolic pathways mediating the activation and restriction of diverse epigenetic ribonucleosides and the consequences of RNA misincorporation of these structures are poorly annotated and understood.

In this study, we investigated a panel of epigenetic pyrimidine and purine ribonucleosides in cultured human cells. We assessed nucleoside cytotoxicity and RNA misincorporation using cell growth assays and RNA mass spectrometry. Furthermore, by employing CRISPR-Cas9 knockouts, we examined the roles of nucleotide salvage enzymes in epigenetic ribonucleoside metabolism. Finally, we evaluated the effects of exogenous ribonucleoside treatment on nucleolar structure and protein translation. We found that purine analogs were generally more toxic than pyrimidine ribonucleosides and characterized the role of ADAL in the restriction of diverse N^6^-modified adenosine incorporation and cytotoxicity, and the role of uridine cytidine kinase 2 (UCK2) in the metabolic activation of pyrimidines. Together, our work provides new insights into the metabolic fate and cellular impact of epigenetic ribonucleosides.

## RESULTS

### Screening cytotoxicity of epigenetic ribonucleosides

To investigate whether epigenetic ribonucleosides can be metabolized and perturb normal biological processes in mammalian cells, we assembled a panel of 18 biologically abundant modified and unmodified ribonucleosides, comprising 8 purines and 10 pyrimidines (Figure 1B).

We screened each nucleoside for cytotoxicity in A549 cells, a human non-small cell lung cancer (NSCLC) line, by adding compounds into normal growth medium at 100 µM concentration and evaluating cell proliferation after 72 h. Among the canonical ribonucleosides (A, G, C, and U), only G significantly inhibited cell proliferation, reducing cell growth by 56.2% ± 3.24% at 100µM (Figure 1C), consistent with a previous report^5^. None of the epigenetic ribopyrimidines tested showed measurable cytotoxicity at 100 µM. Among the modified ribopurines, N^7^- methylguanosine (m^7^G), N^1^-methyladenosine (m^1^A), and inosine (I) were non-toxic. In contrast, bulky N^6^-modified adenosines such as N^6^-isopentenyladenosine (i^6^A) and N^6^, N^6^- dimethyladenosine (m^6,6^A), exhibited strong cytotoxic effects (Figure 1C). We further evaluated the cytotoxicity of i^6^A and m^6,6^A in A549 cells and found that these compounds remained cytotoxic at 10 µM concentration (Supplementary Figure 1A and 1B), aligning with previous reports in diverse human cell lines^19,21,22^. Interestingly, treatment with the structurally related ribonucleoside N^6^-methyladenosine (m^6^A) did not significantly affect the viability of A549 cells under the conditions tested (Figure 1C).

### Bulky N^6^-modified adenosines exhibit cytotoxicity and RNA incorporation

Given the differences observed among various N^6^-modified adenosines, we expanded our panel to include synthetic adenosine analogs with diverse N^6^-substituents (Figure 1D): N^6^- ethyladenosine (e^6^A)^23^, N^6^-cyclopentyladenosine (CPA)^22^, and N^6^-benzyladenosine (Bz^6^A)^22^. We then tested these analogs together with epigenetic N^6^-modified adenosines at 10 µM and 100 µM concentration for cytotoxicity across multiple (primarily cancer-derived) cell lines, including K562, HeLa, U-2 OS, HEK293T, HepG2, MCF7, A549 and MDA-MB-231. A consistent pattern emerged across most cell lines: m^6,6^A, i^6^A and Bz^6^A exhibited strong cytotoxicity, reducing cell viability by more than 80% at 100µM (Figure 1E and Supplementary Figure 1C-J), whereas m^6^A, e^6^A, and CPA were generally non-toxic. We did observe some differences in nucleoside sensitivity across cell lines. For example, HeLa cells were significantly less sensitive to all assayed N^6^-modified adenosines (Figure 1E and Supplementary Figure 1C-J), whereas HepG2 and K562 were most sensitive to m^6,6^A, i^6^A, and Bz^6^A. In addition, m^6^A demonstrated modest toxicity in HEK293T and K562. These differences may stem from variation in metabolic flux or signaling pathways related to nucleic acid quality control or cellular stress across cell types^24,25^.

To investigate whether N^6^-modfied nucleosides are metabolized and incorporated into cellular RNA, we performed quantitative nucleoside LC-QQQ-MS after nucleoside feeding in A549 cells (Supplementary Figure 2 and 3). We could not detect changes in m^6,6^A or m^6^A levels after nucleoside feeding, however the high endogenous abundance of these modifications in total RNA (∼500-1000 ppm) precludes reliable measurement of low-level metabolic incorporation. In contrast, we measured clear incorporation of i^6^A into total RNA, rising by 830 ± 99.0 ppm, a ∼2- fold increase over endogenous levels (Figure 1F). We measured similar i^6^A incorporation in HeLa cells, although it required higher treatment concentration than in A549, in line with the reduced cytotoxicity of i^6^A in HeLa (Supplementary Figure 4A and Figure 1E). For synthetic adenosine analogs, we detected low-level RNA incorporation for e^6^A (7.3 ± 1.4 ppm) and CPA (22.7 ± 0.5 ppm), and higher RNA incorporation for Bz^6^A (150.9 ± 4.3 ppm) (Figure 1F). These results indicate that the cytotoxicity of N^6^-modified adenosine analogs is correlated with the extent of their incorporation into RNA. Interestingly, bulkier N^6^-modifications, such as those found in Bz^6^A and i^6^A, are incorporated more efficiently into RNA than N^6^-modified adenosines with small alkyl substituents. This is in contrast with typical metabolic incorporation trends that favor metabolic activation of nucleotide precursors that are more similar in size to canonical structures^26^, presumably due to increased substrate compatibility with nucleotide salvage enzymes and RNA polymerases.

### ADK activates N^6^-modified adenosine analogs

To understand the metabolic pathways mediating modified adenosine activation and RNA incorporation, we investigated genes in the adenosine salvage pathway. Adenosine nucleosides can be catabolized by adenosine deaminase (ADA) followed by purine nucleoside phosphorylase (PNPase), or phosphorylated by adenosine kinase (ADK) to adenosine monophosphate (AMP), which is then further metabolized to ATP, or alternatively deaminated by AMP deaminase (AMPD) to inosine monophosphate (IMP) (Figure 2A). To test whether ADK mediates the cytotoxic effects of N^6^-modified adenosines, we generated a population knockout (KO) of ADK in A549 cells (Supplementary Figure 5) using CRISPR/Cas9 methods. Strikingly, depletion of ADK resulted in near complete reversal of cytotoxicity induced by N^6^-modified nucleosides, including e^6^A, m^6,6^A, i^6^A and Bz^6^A (Figure 2B). Consistent with this finding, we observed a dramatic 80.9% ± 14.7% reduction of i^6^A and a 98.9% ± 2.0% reduction of Bz^6^A in RNA incorporation in ADK KO cells compared with WT control (Figure 2C, Supplementary Figure 4B). Together, these data demonstrate that ADK is required for RNA incorporation and associated cytotoxicity of N^6^-modified adenosine analogs, likely by mediating their direct phosphorylation to N^6^-modified AMP nucleotides. This metabolic activation likely underlies their cytotoxic effects, further supporting a direct link between RNA incorporation and cytotoxicity.

**Figure 2.**
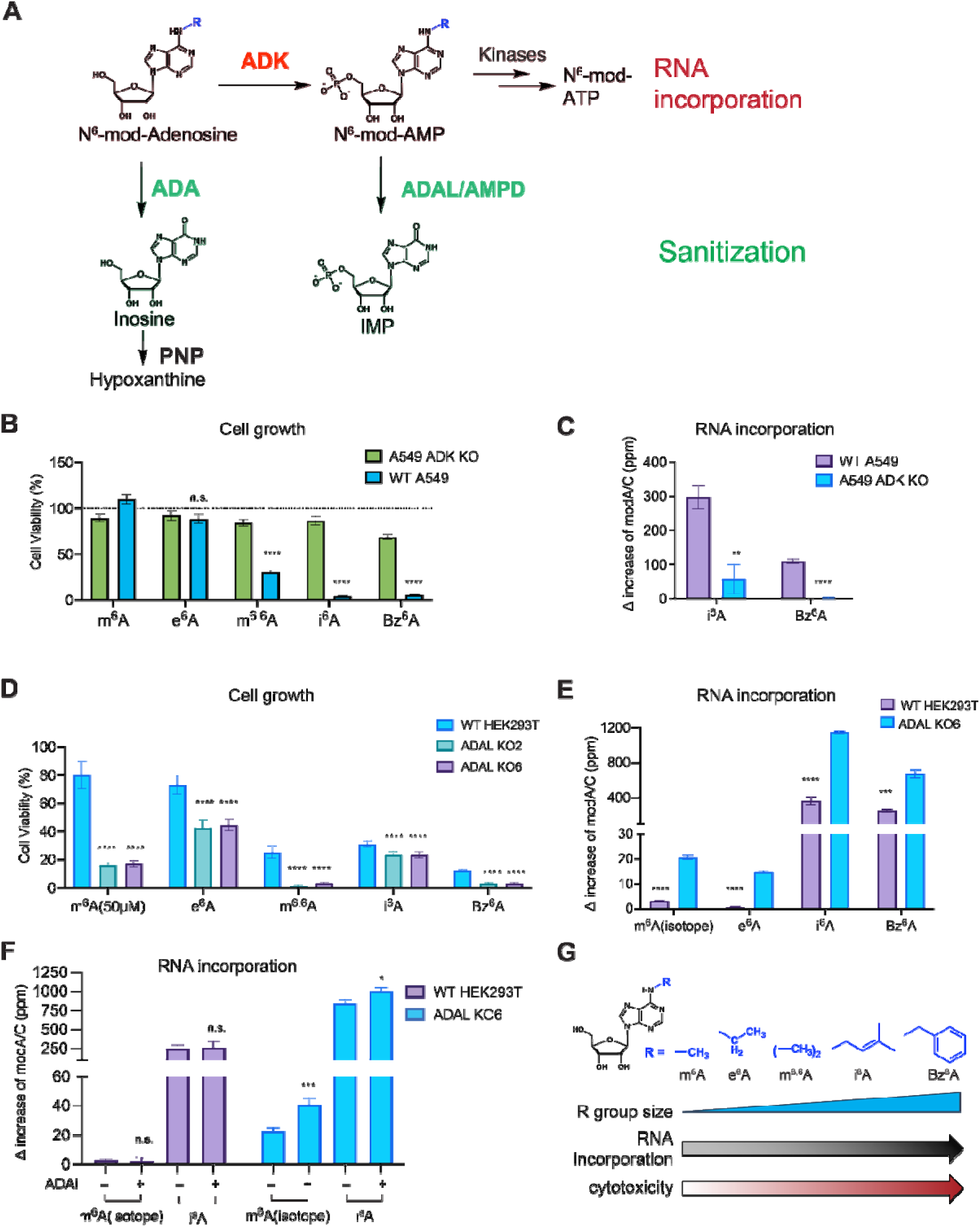
Metabolism of N^6^-modified adenosine analogs. **(A)** Enzymatic pathways involved in adenosine nucleoside metabolism. **(B)** Quantification of cell growth for A549 WT and ADK KO cells after treatment with N^6^-modified adenosines (100 µM for 72 h). Cell growth was measured using an MTS-based assay. Plot represents mean normalized cell viability ± s. d. (*n=9*; three independent biological replicates with three technical replicates for each). Multiple unpaired t-test were performed between WT and ADK KO. Adjusted p values for m^6^A: p<0.000001; for e^6^A: p=0.12; for m^6,6^A: p< 0.000001; for i^6^A: p<0.000001; for Bz^6^A: p<0.000001. **(C)** RNA incorporation for i^6^A and Bz^6^A in A549 WT and ADK KO cells. Cells were treated with 10 µM nucleoside for 24 h and RNA modification levels in total RNA were analyzed by nucleoside LC-QQQ-MS. Data are mean ± s.d. (*n*=3). The unnormalized data was shown in Supplementary Figure 4B. Multiple unpaired t-test were performed between WT and ADK KO. Adjusted p values for i^6^A: 0.001651; for Bz^6^A: 0.000014. **(D)** Quantification of cell growth for HEK293T WT and ADAL KO2/KO6 cells after treatment with N^6^-modified adenosine analogs (10 µM for 72 h for all analogs except m^6^A, which was performed at 50 µM for 72 h). Cell growth was measured using an MTS-based assay. Plot represents mean normalized cell viability ± s. d. (*n=9*; three independent biological replicates with three technical replicates for each). Multiple unpaired t-test were performed between WT and ADAL KOs. Adjusted p values for m^6^A: WT vs KO2 and WT vs KO6: p<0.000001; for e^6^A: WT vs KO2 p<0.000001; WT vs KO6 p<0.000001; for m^6,6^A: WT vs KO2 p<0.000001; WT vs KO6 p<0.000001; for i^6^A: WT vs KO2 p=0.000008; WT vs KO6 p=0.000003; for Bz^6^A: WT vs KO2 p<0.000001; WT vs KO6 p<0.000001. **(E)** RNA incorporation for N^6^-modified adenosine analogs in HEK293T WT and ADAL KO6. Cells were treated with nucleosides (10 µM for 24 h) and modified nucleotide levels in total RNA were analyzed by nucleoside LC-QQQ-MS. Data are mean ± s.d. (*n*=3). The unnormalized data was shown in Supplementary Figure 4C. Multiple unpaired t-test were performed between WT and ADAL KO6. Adjusted p values for m^6^A-D_3_: p=0.000008; for e^6^A: p=0.000003; for i^6^A: p=0.000018; for Bz^6^A: p=0.000117. **(F)** RNA incorporation of m^6^A-D_3_ and i^6^A in HEK293T WT and ADAL KO6 with or without ADAi treatment (10 µM pentostatin). ADAi treated cells were pre-treated with 10µM pentostatin for 6 hours and then exposed to both nucleosides (10 µM) and pentostatin overnight. Cells without ADAi treatment were only incubated with 10µM nucleosides overnight and harvested together with the treated samples. Modified nucleotide levels in total RNA were quantified by nucleoside LC-QQQ-MS. Data are mean ± s.d. Three independent biological replicates were assayed for i^6^A treated cells. For m^6^A-D_3_ feeding, WT without ADAi: *n*=6; ADAL KO6 without ADAi: *n*=5; WT or ADAL KO6 with ADAi: *n*=3. The unnormalized data was shown in Supplementary Figure 8D-E. Multiple unpaired t-test were performed between ADAi treated and untreated samples. Adjusted p-values for m^6^A-D_3_: p=0.00056; for i^6^A: p=0.03. **(G)** Scheme that summarizes the trend of cytotoxicity and RNA incorporation with modification group size at N^6^ position.

### Sanitization of N^6^-modified adenosines by ADAL and ADA

Since the metabolic incorporation of N^6^-modified adenosines followed an unexpected structural trend (i.e. larger derivatives were incorporated more readily than smaller derivatives), we explored whether selective sanitization pathways may exist for smaller N^6^-modified adenosine nucleosides that closely resemble epigenetic nucleosides (i.e. m^6^A). An active sanitization pathway for N^6^-modified adenosines that preferentially removes smaller nucleotides from the nucleotide pool would explain the inefficient metabolism and RNA incorporation of these compounds. We focused our efforts on adenosine deaminase-like protein (ADAL) (Figure 2A), as previous studies have shown that ADAL can deaminate m^6^AMP and m^6^dAMP and prevent misincorporation of N^6^-methyladenine into DNA^17,18^; whether ADAL prevents the misincorporation of N^6^-modified adenosines into RNA is poorly understood. We investigated the role of ADAL in N^6^-modified adenosine sanitization by generating clonal ADAL KO cell lines in HEK293T and HeLa (Supplementary Figure 6A-C, Supplementary Figure 7A and 7B). ADAL KO cells showed no growth defects (Supplementary Figure 6D) and no disruption in endogenous levels of m^6^A, m^6,6^A, or i^6^A (Supplementary Figure 6E). However, deletion of ADAL sensitized cells to all five N^6^-modified adenosine analogs tested, as observed in two independent HEK293T ADAL KO lines and two independent HeLa KO lines (Figure 2D and Supplementary Figure 7C). As the effects were more pronounced in HEK293T, we further focused on these cell lines. Consistent with increased cytotoxicity, ADAL KO also led to increased RNA incorporation of all five analogs (Figure 2E, Supplementary Figure 4C and 4D) with 7-fold or more increase over WT for m^6^A-D_3_ (+17.5 ± 0.8 ppm) and e^6^A (+14.0 ± 0.5ppm), and ∼3-fold increase over WT for i^6^A (+782 ± 47.6 ppm) and Bz^6^A (+420 ± 48.6ppm). We also found that m^6^A-D_3_ incorporation in poly(A) RNA was comparable to that in total RNA (primarily rRNA/tRNA) in ADAL KO cells (Supplementary Figure 4E), suggesting that it can be incorporated into RNA by different polymerases.

Consistent with previous reports showing that ADAL prevents misincorporation of ribonucleotide m^6^A into genomic DNA^17,18^, we measured elevated levels of m^6^dA in our ADAL KO cells that were not fed exogenous m^6^A ribonucleoside (Supplementary Figure 6F). Interestingly, we were not able to measure increased m^6^dA levels in the genome of WT or ADAL KO cells upon feeding exogenous isotopically labeled m^6^A-D_3_ at 50 µM for 2 days (data not shown); we speculate that longer-term feeding or higher nucleoside concentrations would be required to see perturbation of genomic m^6^dA levels; indeed previous studies^18^ used m^6^A-D_3_ for prolonged incubation time to generate detectable levels in genomic DNA.

Whereas ADAL KO sensitized cells to diverse N^6^-modified adenosines and concomitantly increased their incorporation into RNA, incorporation of small alkyl-substituted m^6^A and e^6^A into RNA still proceeded ∼30-70-fold less efficiently than larger i^6^A and Bz^6^A derivatives in the ADAL KO lines. We therefore explored whether sanitization pathways upstream of ADAL, such as deamination by nucleoside deaminases (i.e. ADA1, ADA2)^28^, may prevent the metabolism of small N^6^-modified nucleosides. Notably, ADA1 shares a high degree of structurally similarity with ADAL, with a root-mean-square-deviation (RMSD) of only 1.56 Å (Supplementary Figure 8A-C). To inhibit the activity of ADA proteins, we treated WT and ADAL KO cells with pentostatin, an established and potent small molecule inhibitor of ADA1/2 enzymes^29,30^, and measured RNA incorporation of m^6^A-D_3_ and i^6^A nucleosides. Pentostatin treatment modestly increased RNA incorporation of m^6^A-D_3_ in ADAL KO cells (82% increase), but not in WT cells, and had minimal effect on incorporation of i^6^A (19% increase in KO) (Figure 2F, Supplementary Figure 8D and 8E). Collectively, our results indicate that diverse N^6^-modified adenosine analogs are activated by ADK and incorporated into RNA. Bulky N^6^-modified adenosines exhibit the highest levels of cytoxicity and RNA incorporation, and cells can mitigate misincorporation and cytotoxicity of N^6^- modified adenosine analogs through sanitization pathways involving nucleotide and nucleoside deaminases, ADAL and ADA (Figure 2G); however, even in the absence of ADAL/ADA-mediated sanitization pathways, m^6^A and structurally related derivatives show limited RNA misincorporation, suggesting that other mechanisms exist to prevent stochastic RNA incorporation of epigenetic N^6^-modified adenosines.

### Cytotoxicity of epigenetic pyrimidine ribonucleosides

Although epigenetic ribopyrimidines did not show cytotoxic effects in our initial screen at 100 µM concentration in A549 cells (Figure 1C), we repeated the screen at higher concentrations and expanded to additional human cell lines. In A549, we began to see cytotoxic effects at 5 mM concentration (Figure 3B, Supplementary Figure 9A). Among the compounds tested, dihydrouridine (D) inhibited cell growth by 83.7% ± 1.3%, 5-methyluridine (m^5^U) by 50.3% ± 4.0%, 5-methylcytidine (m^5^C) by over 90%, and 2’-O-methylcytidine (Cm) by 63.8% ± 2.6% (Figure 3B). These results indicate that epigenetic ribopyrimidines can be cytotoxic, but only at high (millimolar) concentrations. We then extended the assay to a panel of cell lines. At 1 mM treatment, m^5^C consistently exhibited the strongest cytotoxicity, inhibiting growth by over 90% in HeLa and HEK293T cells (Figure 3C, Supplementary Figure 9C-H). Interestingly, pseudouridine (Ψ) showed cell-type specific effects, with more than 90% inhibition in HeLa cells, but minimal toxicity in A549 and MCF7. Moderate inhibition (∼30%) was observed in HepG2, U2OS and HEK293T cells. When the treatment concentration was further increased to 5 mM, the cytotoxicity patterns become more pronounced: D, m^5^C and m^5^U were broadly toxic across most cell lines, while Ψ remained selectively toxic in specific cell types (Supplementary Figure 9B-H).

**Figure 3.**
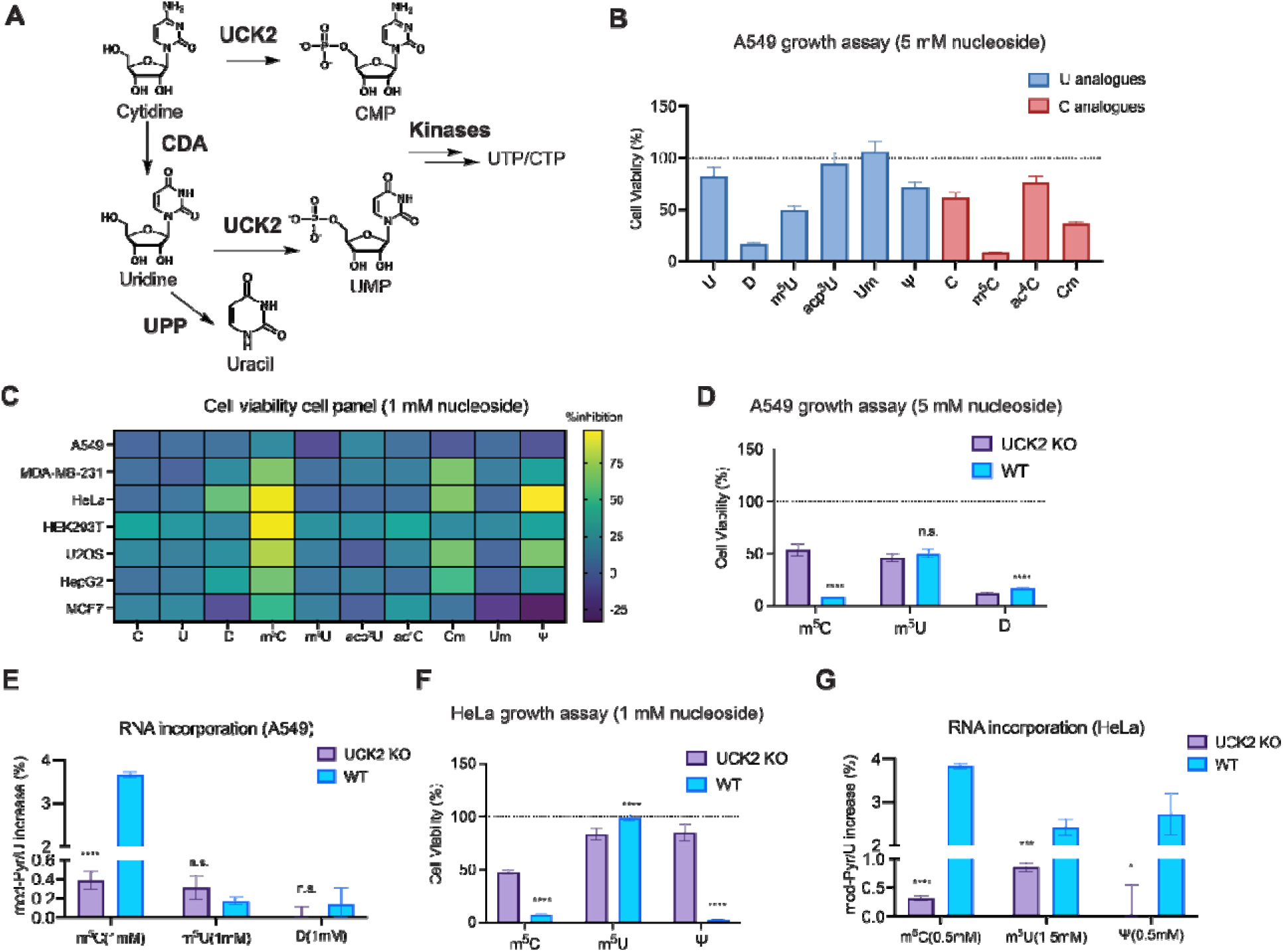
Metabolism of modified ribopyrimidine analogs. **(A)** Metabolic pathways involved in ribopyrimidine metabolism. **(B)** Cytotoxicity of epigenetic ribonucleosides (5 mM for 72 h) in WT A549 cells. Cell viability was measured by MTS assay and plot represents mean normalized cell viability ± s. d. (*n=9*; three independent biological replicates with three technical replicates for each). **(C)** Ribopyrimidine nucleoside cytotoxicity in a panel of human cell lines. Heatmap shows percentage growth inhibition (1 mM nucleoside for 72 h), corresponding to Supplementary Figure 9A and 9C-H. Cell viability was measured by an MTS-based assay and represents mean normalized by untreated cells. (*n*=9 with three technical replicates for each of the three independent biological replicates). **(D)** Quantification of cell growth for A549 WT and UCK2 KO cells with m^5^C, m^5^U or D treatment. Cell growth was measured by MTS assay after nucleoside treatment (5 mM for 72 h). Plot represents mean normalized cell viability ± s. d. (*n=9*; three independent biological replicates with three technical replicates for each). Multiple unpaired t-test were performed between WT and UCK2 KO. Adjusted p values for m^5^C: p<0.000001; for D: p<0.000001. **(E)** RNA incorporation levels for m^5^C, m^5^U or D. A549 WT or UCK2 KO cells were treated with 1 mM nucleoside for 24 h and RNA modification levels in total RNA were quantified by nucleoside LC-QQQ-MS. Data are mean ± s.d. (*n*=3). The unnormalized data was shown in Supplementary Figure 10E. Multiple unpaired t-test were performed between WT and UCK2 KO. Adjusted p value for m^5^C: p=0.000004. **(F)** Quantification of cell growth for HeLa WT and UCK2 KO cells after treatment with m^5^C, m^5^U or Ψ (1 mM for 72 h). Cell growth was measured by MTS assay. Plot represents mean normalized cell viability ± s. d. (*n=9*; three independent biological replicates with three technical replicates for each). Multiple unpaired t- test were performed between WT and UCK2 KO. Adjusted p values for m^5^C: p<0.000001; for m^5^U: p=0.000009; for Ψ p<0.000001. **(G)** RNA incorporation levels for m^5^C, m^5^U or Ψ in HeLa WT or UCK2 KO cells. Modified nucleotide levels were quantified in total RNA by nucleoside LC-QQQ-MS after m^5^C/ Ψ (0.5 mM) or m^5^U (1.5 mM) treatment for 24 h. Data are mean ± s.d. (*n*=3). Unnormalized data is hown in Supplementary Figure 11D. Multiple unpaired t-test were performed between WT and UCK2 KO. Adjusted p values for m^5^C: p<0.000001; for m^5^U: p=0.00045; for Ψ: p=0.018.

### UCK2 mediates the incorporation and toxicity of epigenetic ribopyrimidines

Our group^31–33^ and others^34–37^ have previously established that uridine-cytidine kinase 2 (UCK2), which phosphorylates uridine/cytidine nucleosides to NMPs, is rate-limiting for uptake and RNA incorporation of modified pyrimidine ribonucleosides (Figure 3A). To explore its role in metabolism of epigenetic ribopyrimidines, we generated UCK2 KO cells in A549 (Supplementary Figure 10A) and HeLa (Supplementary Figure 11A) and assayed sensitivity to D, m^5^C and m^5^U. In A549, UCK2 depletion markedly rescued the cytotoxicity induced by m^5^C, while no significant rescue was observed for D or m^5^U treatments (Figure 3D, Supplementary Figure 10B and 10C). Through dose-response titrations, we observed >7-fold increase in the IC_50_ of m^5^C in A549 cells with UCK2 depletion. In contrast, the IC_50_ values for m^5^U and D remained nearly unchanged between UCK2 KO and WT cells (Supplementary Figure 10D). Using nucleoside LC-MS/MS, we quantified RNA incorporation (Supplementary Figure 12) and detected 3.65% ± 0.07% incorporation of m^5^C in WT A549 cells (a 4-fold increase over endogenous m^5^C levels), compared to only 0.38% ± 0.096% incorporation in UCK2 KO cells (Figure 3E, Supplementary Figure 10E). In contrast, D and m^5^U levels did not significantly change between UCK2 KO and WT (Figure 3E, Supplementary Figure 10E), consistent with similar cytotoxicity profiles (Figure 3D).

Similar trends were observed in HeLa cells. UCK2 KO rescued m^5^C-induced toxicity but not m^5^U toxicity (Figure 3F, Supplementary Figure 11B), suggesting m^5^U is a poor substrate for UCK2. As noted above, HeLa cells were selectively toxic to Ψ, an effect that we found to be dependent on UCK2 status (Figure 3F and Supplementary Figure 11B and 11C). This effect was less pronounced but still evident in A549 cells (Supplementary Figure 10C). IC_50_ values for m^5^C and Ψ in HeLa UCK2 KO cells increased ∼4-5-fold as compared to WT cells (Supplementary Figure 11C. In WT HeLa cells, LC-QQQ-MS quantification revealed UCK2-dependent incorporation of m^5^C at 3.83% ± 0.07% and Ψ at 2.72% ± 0.49% following 0.5 mM treatment in WT HeLa (Figure 3G, Supplementary Figure 11D). Surprisingly, although m^5^U did not show UCK2-dependent toxicity in HeLa, we detected UCK2-dependent RNA incorporation: m^5^U levels increased by ∼14.5-fold in WT (2.41% ± 0.20% incorporation) and by only 5-fold in UCK2 KO (0.86% ± 0.08% incorporation) (Figure 3G, Supplementary Figure 11D). This suggests that m^5^U toxicity may not be directly related to metabolism or RNA incorporation. Taken together, these results demonstrate that UCK2 is the key enzyme responsible for m^5^C and Ψ toxicity via its role in promoting their RNA incorporation. In contrast, m^5^U and D show minimal UCK2 dependence, suggesting distinct metabolism or cytotoxicity stemming from the unprocessed nucleosides.

### Pyrimidine ribonucleoside cytotoxicity induces DNA damage but is not associated with DNA incorporation

We next investigated the mechanisms by which modified ribonucleosides cause cell death. Given the high concentrations of epigenetic pyrimidine ribonucleosides needed for cytotoxicity, and associated high levels of RNA incorporation, we questioned whether modified pyrimidines fed in this way may also be metabolized to modified deoxynucleotides and possibly incorporated into DNA, thereby inducing DNA damage or replication stress/inhibition. First, we investigated whether modified ribonucleoside treatment induced DNA damage in A549 or HeLa cells. In HeLa cells, treatment with m^5^C, Ψ, or m^5^U for 48h at concentrations that induced comparable RNA incorporation (∼3%) resulted in DNA damage as measured by γH2AX Western blot, whereas i^6^A treatment did not increase γH2AX signal in either cell line (Supplementary Figure 13A and 13B). In A549, γH2AX induction was more modest, however m^5^C treatment did result in a slight increase in DNA damage-associated signal. We next focused on quantifying DNA incorporation and deoxynucleotide metabolites derived from m^5^C as it was the most toxic epigenetic ribopyrimidine we assayed, it induced DNA damage in HeLa and A549 cells, and due to the availability of corresponding modified deoxyribonucleoside standards for LC-MS analysis. We extracted genomic DNA from treated and untreated cells and analyzed 5-methyl-2’- deoxycytidine (5mdC) levels by nucleoside LC-QQQ-MS. Upon m^5^C feeding, we did not observe significant differences in 5mdC levels in either A549 or HeLa cells (Supplementary Figure 13C and 13D). However, considering the high endogenous abundance of 5mdC in genomic DNA, it is possible that subtle changes might be masked in bulk LC-QQQ-MS analysis. Further, we examined the nucleotide pools and found no detectable formation of m^5^dCTP after 2 h or 24 h m^5^C treatments (Supplementary Figure 14). Instead, we observed a marked accumulation of m^5^C monophosphate (m^5^CMP) and m^5^C triphosphate (m^5^CTP), as well as the corresponding deamination products, m^5^U monophosphate (m^5^UMP) and m^5^U triphosphate (m^5^UTP) (Supplementary Figure 14), aligning with increases in m^5^C in RNA (Figure 3G, Supplementary Figure 11D). The abundance of m^5^C and m^5^U-derived nucleotide was higher 2 h post-feeding than 24 h, indicating that cells were actively metabolizing these molecules. In contrast, whereas m^5^C nucleoside levels also declined from 2 h to 24 h, m^5^U nucleoside and 5-methyluracil (thymine) accumulated in the cell from 2 h to 24 h. Taken together, our findings show that epigenetic ribopyrimidine-induced cytotoxicity correlates with DNA damage, but appears to be independent of metabolism to modified deoxynucleotides or DNA incorporation.

### i^6^A and **Ψ** induce nucleolar stress through distinct mechanisms

Active rDNA transcription is essential for nucleolar organization, and nucleolar function is tightly linked to its structure as a multiphase condensate^38–41^. The mammalian nucleolus consists of three subcompartments: the fibrillar center (FC), dense fibrillar component (DFC) and granular component (GC). rDNA is transcribed at the FC-DFC interface, pre-rRNA processing occurs in the DFC, and ribosomal subunit assembly takes place in the GC^38–42^. Inhibition of rDNA transcription triggers nucleolar stress, characterized by redistribution of nucleolar proteins to the periphery, leading to a rounded, condensed appearance^39–42^. This can be visualized using markers specific to different nucleolar compartments^40,42^.

To examine whether epigenetic ribonucleoside cytotoxicity and RNA incorporation correlate with perturbation of nucleolar structure and function, which could lead to defects in ribosome biogenesis and protein translation and arrest cell cycle progression, we examined nucleolar morphology by staining for nucleophosmin (NPM1), a GC maker, and RPA-194 (RNA Polymerase I Subunit A), an FC marker. In HeLa cells treated with i^6^A treatment, we observed nucleolar caps forming around the GC (Figure 4A), a hallmark of nucleolar segregation caused by Pol I inhibition^43,44^. Quantification of nucleolar eccentricity showed a 27% decrease (Figure 4B), reflecting nucleolar rounding in the images (Figure 4A) and consistent with Pol I inhibition^45,46^. In addition, NPM1 partitioning was significantly altered by i^6^A treatment (Figure 4C). While Ψ treatment did not induce nucleolar caps, it did cause nucleolar rounding, as evidenced by a 14% decrease in eccentricity (Figure 4A and 4B). These morphological changes suggest that Ψ also induces nucleolar stress, albeit likely through a mechanism distinct from Pol I inhibition. In contrast, m^5^C treatment had no obvious effect on nucleolar morphology (Figure 4A and 4B).

**Figure 4.**
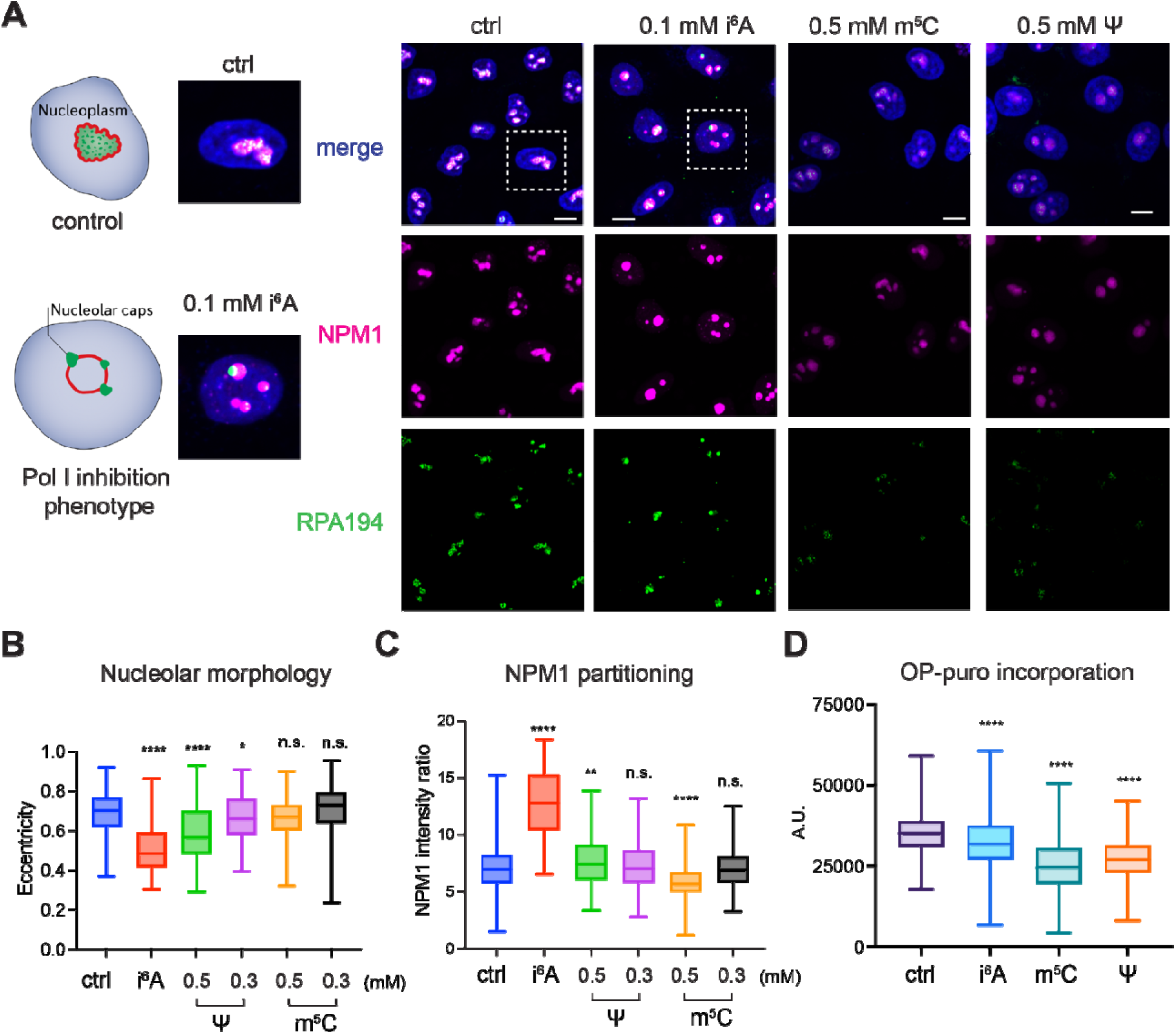
Epigenetic nucleosides induce nucleolar stress and global translation inhibition. **(A)** Confocal micrographs of HeLa cells treated with 100 µM i^6^A, 500 µM m^5^C or 500 µM Ψ for 24h. Cells were stained with antibodies for NPM1 and RPA-194 and a nuclear dye. Scale bar represents 10 µm. Representative images from two biological replicates. **(B)** Quantification of nucleolus morphology (eccentricity). Boxes in the plot represent 25^th^ to 75^th^ percentiles and whiskers show min to max. Two tailed unpaired t-test was performed between treated and control samples and p values for i^6^A: p<0.000001; Ψ (0.5 mM): p=0.000011; Ψ (0.3 mM): p=0.047; 40-120 cells were analyzed per condition. **(C)** Quantification of NPM1 partitioning. Boxes in the plot represent 25^th^ to 75^th^ percentiles and whiskers show min to max. Two tailed unpaired t-test was performed between treated and control samples and p values for i^6^A: p<0.000001; Ψ (0.5 mM): p=0.0062; m^5^C: p<0.000001. Between 40-300 cells were analyzed per condition. **(D)** Global translation levels upon nucleoside feeding. Quantification of OP-puro incorporation (Cy3 signal) from HeLa cells treated with 100 µM i^6^A, 500 µM m^5^C, or 500 µM Ψ for 24h. Boxes in the plot represent 25^th^ to 75^th^ percentiles and whiskers show min to max. Two tailed unpaired t-test were performed between treated and control samples and p values are for i^6^A: p<0.000001; for m^5^C: p<0.000001; for Ψ: p<0.000001.

We observed similar nucleolar perturbation with i^6^A treatment in A549 cells (Supplementary Figure 15A), though changes in eccentricity and NPM1 partitioning were less pronounced (Supplementary Figure 15B and 15C). These differences likely reflect cell-type-specific responses. Nonetheless, the consistent formation of nucleolar caps upon i^6^A treatment in both HeLa and A549 cells supports its role in Pol I inhibition. In contrast, m^5^C treatment did not induce noticeable alterations in A549 cells (Supplementary Figure 15), consistent with observations in HeLa cells (Figure 4A). Together, these findings suggest that i^6^A inhibits cell growth by inhibiting Pol I and inducing nucleolar stress, presumably by first misincorporating into rRNA. Ψ also induces nucleolar stress, possibly by interfering with ribosome biogenesis at the level of pre-rRNA processing or modification, given the role of pseudouridylation in these processes^47^. The absence of effects with m^5^C suggests its cytotoxicity operates through mechanisms unrelated to nucleolar function.

### Epigenetic nucleosides inhibit global translation

Finally, we employed *O*-propargyl-puromycin (OP-puro)^48^, which incorporates into nascent polypeptides, to assess global protein synthesis after epigenetic nucleoside treatment. We observed a marked reduction in OP-puro signal following epigenetic nucleoside treatments: 27.7% ± 1.4% (mean ± SEM) decrease for m^5^C, 22.9% ± 1.2% (mean ± SEM) decrease for Ψ, and 8.6% ± 1.4% (mean ± SEM) decrease for i^6^A (Figure 4D, Supplementary Figure 16). These results indicated that all three epigenetic nucleosides disrupt global translation to varying extents.

## DISCUSSION

In this study, we show that a subset of epigenetic ribonucleoside can be metabolized by cells and incorporated into RNA, with associated cytotoxicity. We found that N^6^-modified adenosines were the most toxic of those tested, and further showed that their metabolic activation relies upon ADK and is opposed by ADAL- and ADA-mediated sanitization pathways. Using a broader panel of epigenetic and synthetic N^6^-modified adenosines, we found a strong restriction against the misincorporation of adenosine analogs with small N^6^-alkyl substitutions such as the abundant epigenetic ribonucleoside m^6^A. Compared to purine analogs, pyrimidines were less toxic but demonstrated measurable UCK2-dependent and UCK2-independent cytotoxicity at high concentrations in multiple cell lines. Furthermore, we show that i^6^A and Ψ misincorporation correlates with nucleolar stress and global translation inhibition.

Our findings on the role of ADAL in deaminating N^6^-methyladenosine nucleotides align with previous work in *Arabidopsis* and human cells, where ADAL were shown to serve a similar protective function^16^. In addition, we show that ADAL has broad substrate specificity and can process a variety of native and synthetic N^6^-modified adenosines including i^6^A, Bz^6^A, and e^6^A, and prevent their misincorporation into RNA. Similarly, ADK possesses broad substrate specificity and can activate diversely functionalized N^6^-modified. While this manuscript was in preparation, Ogawa *et al.* reported in a preprint that m^6^A, m^6,6^A and i^6^A can be phosphorylated by ADK and deaminated by ADAL^49^, corroborating our findings. By measuring RNA misincorporation rates of different N^6^-modified adenosines, we found that larger N^6^-substituents, such as isopentenyl and benzyl, enabled incorporation at >100-fold higher levels than small alkyl N^6^-substitutions (i.e. methyl and ethyl) (Figure 2E). This is in stark contrast to typical metabolic incorporation trends for modified nucleosides that favor activation and incorporation of nucleosides that more closely resemble canonical nucleoside structures. One possible explanation is that ADAL more readily accommodates NMPs with small N^6^-modifications, as suggested by a structure–activity relationship (SAR) study^50^, presumably as this enzyme has evolved primarily to process m^6^A nucleotides due to its abundance. However, even in ADAL KO cells, larger N^6^-modified adenosines still incorporate into RNA at >100-fold higher levels than m^6^A or e^6^A, suggesting that ADAL is not the only (and perhaps not even the major) restriction mechanism for preventing m^6^A misincorporation. We show that adenosine deaminase (ADA) serves a minor role in sanitizing exogenous m^6^A nucleosides. What other pathways may function to restrict unwanted m^6^A incorporation in RNA (and DNA)? In addition to other enzymes that sanitize the nucleotide pool, differential uptake into cells could be mediated by nucleoside transporters. Further, RNA misincorporation could be managed by RNA polymerases or by enzymes that can remove unwanted m^6^A nucleotides after RNA synthesis.

In contrast to purine ribonucleosides, pyrimidines (both modified and unmodified) did not exhibit cytotoxicity except at millimolar concentrations. At these high concentrations, cytotoxicity of epigenetic pyrimidine ribonucleosides largely depended upon UCK2, which aligns with previous studies implicating UCK2 as the rate-limiting enzyme for salvage of cytotoxic pyrimidine ribonucleoside analogs^31–37^. UCK2 is known to have stringent substrate selectivity^32,51^ and therefore this kinase (and NMP kinase CMPK1) may serve as a filter to prevent metabolism and subsequent RNA misincorporation of epigenetic pyrimidines. Whereas we measured high levels of RNA misincorporation correlating with cytotoxicity, it is unknown whether cells tolerate lower concentrations of these nucleosides due to inefficient metabolism (or sanitization pathways) or lack of major perturbation to RNA structure/function upon misincorporation of pyrimidine modifications. Catabolic enzymes for Ψ (which is the most abundant RNA modification) have been reported in bacteria^52^ and in plants^53^, but homologous pathways in mammals are not known.

We evaluated the cytotoxicity mechanisms of epigenetic ribonucleosides by characterizing nucleolar morphology and measuring protein translation. Several synthetic nucleosides and nucleobases are known to induce nucleolar stress, including 5-fluorouracil^54^, 5-fluorouridine^54^ and 4-thiouridine^55^. Additionally, DHODH inhibitors have been shown to disrupt nucleotide balance and trigger nucleolar rounding^56^. In our studies, modified epigenetic ribonucleosides can similarly perturb nucleolar function, but through diverse mechanisms. Treatment with i^6^A induces nucleolar caps, consistent with Pol I inhibition. Yakita *et al.* also demonstrated that i^6^A inhibits Pol I transcription by RT-qPCR analysis of rRNA transcripts^19^. Since i^6^A also misincorporates into RNA, we speculate that it may serve as a transcription terminator after accumulating in rRNA. In contrast, Ψ induces nucleolar stress without significant nucleolar cap formation, suggesting that Ψ misincorporation perturbs later steps of rRNA processing and ribosome biogenesis^47^, rather than inhibiting Pol I transcription. In *E. coli*, excessive Ψ in rRNA has been shown to inhibit ribosome assembly^57^. Whereas m^5^C did not induce apparent nucleolar stress based on FC and GC morphology, it did inhibit protein translation (as did i^6^A and Ψ). Therefore, m^5^C may act at later stages of ribosome biogenesis or translation inhibition may be due to mRNA misincorporation; indeed, m^5^C modifications in synthetic mRNA are known to perturb translation efficiency^58^.

In summary, our study provides new insights into the cellular metabolism of epigenetic ribonucleosides and highlights the role of nucleotide salvage and sanitization pathways in managing their incorporation into RNA. These findings deepen our understanding of the metabolic fate of RNA modifications following RNA turnover and offer valuable insights into the cellular handling of nucleoside-based drugs with similar structures.

## ACKNOWLEDGEMENTS

We thank John Eng, Venu Vendavasi and Christina J. DeCoste at the Princeton Mass Spectrometry, Biophysics and Flow Cytometry core facilities, respectively, for providing support for nucleoside LC-MS, plate reader-based assays and flow cytometry. R.E.K. acknowledges support from the Ludwig Princeton Branch, NIH (R01 GM132189) and NSF (MCB-1942565).

J.D.R. and J.A.B. acknowledge support from the Ludwig Princeton Branch. X.S. was supported by a generous gift from the Edward C. Taylor 3rd Year Graduate Fellowship in Chemistry.

