## Supplementary Information for "Metabolism of epigenetic ribonucleosides leads to nucleolar stress and cytotoxicity"

Table of contents

Methods 2

Supplementary Figures 8

**Methods**

*Nucleosides*

Modified nucleosides were purchased from the following vendors: $m^1A$ (Jena Biosciences, #n1055), $Bz^6A$ (Biosynth, #NB06325), CPA (Biosynth, #NC45738), $e^6A$ (Cayman, #28428), $i^6A$ (Cayman, #20522), $m^6A$ (Ark Pharm, #AK153126), $m^{6,6}A$ (Cayman, #35348), Ψ (Biosynth, #NP11297), $m^5U$ (TCI, #M1405), D (Apollo Sci, #or17694), Um (Biosynth, #42591899), $m^5C$ (Biosynth, #NM03720), $ac^4C$ (Bionet, #as2045), Cm (Biosynth, #nm06302), I (Cayman, #34373), $m^7G$ (Cayman, #15988).

*Cell culture*

HEK293T, A549, HeLa, MDA-MB-231, MCF-7, U2OS, and HepG2 cells were cultured in high glucose DMEM (Life Technologies) supplemented with 10% fetal bovine serum (Atlanta), 1x penicillin-streptomycin (Gibco Life Technologies) and 2 mM L-glutamine (Life Technologies) at 37°C in a humidified atmosphere with 5% CO2 and atmospheric oxygen concentration.

*KO cell line generation*

For selection of UCK2 KO cell lines, HeLa or A549 WT cells ($0.8 \times 10^6$) were seeded in a six-well dish one day before transfection. 2µg pX330-hspCas9 plasmid containing UCK2 sgRNA (TCTGCTCCGAGGTAAGGACA) and pcDNA3-FKBP-EGFP-HOTag3 (Addgene # 106924, 200 ng) were co-transfected using Lipofectamine 3000 (Thermo Scientific) following the manufacturer's instruction. Fresh media was added 24 h post-transfection. The top 10% of cells with green fluorescent protein (GFP) signal were sorted by FACS 48 h after transfection into 96-well plates. For generation of ADAL and ADK KO cell lines, puromycin-based cell selection was used. For ADK populational KO, A549 WT cells ($0.5 \times 10^6$) were seeded in a 6-well plate and the following day, 2µg pX459-pspCas9-Puro plasmid containing ADK sgRNA (ACAGCAGAGATGTCAAGCAG) was transfected with Lipofectamine 3000. After 24 h, cells were selected with 0.5 μg/ml puromycin for 3 days. Cells were then recovered in regular medium for 24 h and ready for assay. For ADAL single-cell-derived KO, HEK293T WT cells (0.1 x$10^6$) were seeded in a 24-well plate. After 24 h, 500 ng pX459-pspCas9-Puro plasmid containing ADAL sgRNA (GGAGACTGTAAAACTTGCCG) was transfected with Lipofectamine 3000. After 24 h, cells were selected with 1 μg/ml puromycin for 3 days. Cells were then recovered in regular medium for 24 h and diluted to single cells into 96-well plates. Surviving colonies were expanded to 6-well plates and then 10-cm plates and KO was confirmed by genomic PCR and western blot.

*Cell viability assay*

Cells were seeded in 96-well Falcon clear plates (2,000 cells in 200 μl of medium per well) 24 h before treatment. Cells were treated with nucleosides at the indicated concentrations. Cell viability was measured using the MTS assay (CellTiter 96 Aqueous Non-Radioactive Cell Proliferation Assay; Promega, G5430) at 72 h post-treatment following the manufacturer's instructions. The absorbance of each well was measured at 490 nm on a Synergy H1 Microplate Reader (BioTek). All treated cells were normalized to the untreated sample. Three independent biological replicates and three technical replicates were collected for each condition. For generating titration curves, normalized cell viability was fitted based on a 4-parameter dose-response equation using GraphPad Prism (version 10.1.1):

$$Y=Bottom + (Top-Bottom)/(1+10^{((logIC50-X) \, x \, HillSlope)}) \qquad (1)$$

*Total RNA isolation and nucleoside LC-QQQ-MS analysis*

Total RNA from treated or untreated cells was extracted using TRIzol reagent (Thermo Fisher) according to the manufacturer's protocol. RNA samples (3 μg) were digested in a 20 μL reaction containing 2 units of Nuclease P1 (Wako, dissolved in H2O) in 7 mM NaOAc and 0.4 mM ZnCl2 at 37°C for 2h. The resulting nucleotides were dephosphorylated at 37°C for 2 h by 2 units of Antarctic Phosphatase (NEB) at a final concentration of 1X Antarctic phosphatase buffer.

Ribonucleoside were quantified using an Agilent 1290 LC Infinity II system coupled to an Agilent 6495C LC/TQ module with a Hypersil Gold aQ column. Both chromatographic separation and mass spectrometric conditions, including the LC gradient and MRM transitions, were adapted from previously reported methods in literature[1,2]. Measurement was taken from at least 3 distinct biological replicates. Relative levels of modified nucleosides were calculated by normalizing the concentration of modified nucleoside to the canonical nucleoside (e.g., C or A). All quantification was processed on Agilent MassHunter Workstation Data Acquisition version 10.0.

*Western blotting*

Adherent cells were harvested by scraping into ice-cold PBS buffer and pelleted. For whole cell lysate, cells were lysed in RIPA buffer (10 mM Tris pH 8.0, 140 mM NaCl, 1 mM EDTA, 0.5 mM EGTA, 1% Triton X-100, 0.1% sodium deoxycholate, 0.1% SDS, 1 mM PMSF). For DNA damage assay, cells were lysed in LB3 buffer (10mM Tris pH 8.0, 100 mM NaCl, 1 mM EDTA, 1 mM EGTA, 0.1% sodium deoxycholate and 0.5% sodium laurylsarcosine with 1 mM PMSF and Roche protease inhibitor tablet) and followed by sonication. Proteins were separated on a 10-15% SDS-PAGE gels first and then transferred to nitrocellulose membranes (VWR). Membranes were blocked with 5% BSA in TBST (5 mM Tris-HCl pH 7.5, 15 mM NaCl, 1% Tween-20) or 5% milk in TBST. Primary antibodies were applied in the following dilutions: UCK2 (Proteintech), 1:1000; β-actin (Cell Signaling), 1:10,000; ADAL (Proteintech), 1:1000; γH2AX (Millipore), 1:1000. After incubation with IR-labeled secondary antibodies and washes with TBST buffer, blots were imaged on a LI-COR Odyssey Imaging System.

*Immunofluorescence*

Cells were seeded onto round glass 12 mm coverslips (Fisher Scientific) at 0.05 x $10^6$ cells per well in a 24-well plate. Cells were allowed to grow overnight and were treated with nucleoside at indicated concentrations for 24 h. Coverslips were washed with PBS twice and cells were fixed for 15 min at 37°C in PBS containing 3% paraformaldehyde (pH 7.3) and washed with PBS twice. Cells were permeabilized with PBST (PBS + 0.1% Triton X-100) for 15 min at room temperature and washed twice with PBS. The coverslips were blocked with 5% goat serum for 1 h at room temperature, then incubated with primary antibody for 2 h at room temperature. Primary antibodies were applied in the following dilutions: RPA194-AF488 (Santa Cruz), 1:400; NPM1-AF647 (Thermo Fisher), 1:200 (incubated for 5h at 37°C); G3BP1 (Santa Cruz), 1:200. After washing three times with PBS for 5 min each, secondary antibody incubations were performed for 1 h at RT in dark or skip the secondary antibody if fluorophore is conjugated to the primary antibody. After three rounds of PBS washes, nuclei were stained with Hoechst 33342 with a 1:10,000 dilution (Thermo) in PBS for 10 min, and coverslips were washed with PBS twice for 5 min. Coverslips were mounted on cell-face down atop a microscope slide with ProLong Gold AntiFade Reagent (Life Technologies). Fixed cells on the microscope slides were imaged on a Nikon Eclipse Ti microscope equipped with a 60x objective, a CMOS camera, and the NIS Elements AR software.

*Confocal microscopy*

Cells were imaged on a Yokogawa CSU-W1 spinning-disc confocal microscope using a Nikon 60X oil-immersion Plan Apo λD objective (numerical aperture = 1.42) on a Nikon Eclipse Ti2 body with an ORCA-Fusion BT back-thinned high quantum efficiency sCMOS camera. DAPI, RPA194, and NPM1 were imaged using 405, 488, and 647 nm laser lines, respectively.

*Image analysis*

Image analysis was done in Cell Profiler software version 4.2.6. First, nuclei were segmented using the DAPI channel and 'Identify Primary Objects' module with global Otsu thresholding and nucleoli were segmented using the NPM1 channel and 'Identify Primary Objects' with adaptive minimum cross-entropy thresholding. Then, the 'Identify Tertiary Objects' module was used to identify nucleoplasm by subtracting she previously thresholded smaller objects (nucleoli) from the previously thresholded larger objects (nuclei). Finally, nucleolar and nuclear morphology features were quantified using the 'Measure Object Size Shape' module and NPM1 intensity was measured in the nucleoplasm and nucleolus using the 'Measure Object Intensity' module. For all morphology and partitioning measurements, nucleoli within a single nucleus were assigned to its nucleus using the 'Relate Objects' module and represent per cell averages. NPM1 partitioning was measured as the ratio of average NPM1 intensity in nucleoli of an individual cell and its corresponding nucleoplasm as described previously[3]. Data was plotted in GraphPad Prism version 10.1.1.

*Global translation assay*

Global translation efficiency was evaluated by treating cells with O-propargyl puromycin (OP puro, Click Chemistry Tools) and detecting fluorescence after click chemistry via fluorescence microscopy following literature precedent[4,5]. For quantification of OP-puro signal, Fiji ImageJ (2.14.0/1.54f) was used to mask cells according to DAPI staining and align with the Cy3 signal to assess intensity within individual cells. The resulting measurement of mean intensity density per area was used to compare translational efficiency across different treatment conditions.

*Nucleotide metabolomics*

Metabolites from cells were extracted in 80% methanol on dry ice and further analyzed using a Vanquish Horizon UHPLC System (Thermo Scientific) coupled to an Orbitrap Exploris 240 Mass Spectrometer (Thermo Scientific). Waters XBridge BEH Amide XP Column (particle size, 2.5 μm; 150 mm (length) × 2.1 mm (i.d.)) was used for hydrophilic interaction chromatography (HILIC) separation. Column temperature was 25 °C. Mobile phases A = 20 mM ammonium acetate and 22.5 mM ammonium hydroxide in 95:5 (v/v) water:acetonitrile (pH 9.45) and B = 100% acetonitrile were used for both ESI positive and negative modes. The linear gradient eluted from 90% B (0.0–2.0 min), 90% B to 75% B (2.0–3.0 min), 75% B (3.0–7.0 min), 75% B to 70% B (7.0–8.0 min), 70% B (8.0–9.0 min), 70% B to 50% B (9.0–10.0 min), 50% B (10.0–12.0 min), 50% B to 25% B (12.0–13.0 min), 25% B (13.0–14.0 min), 25% B to 0.5% B (14.0–16.0 min), 0.5% B (16.0–20.5 min), 90% B (20.5 – 25.0 min). The flow rate was 0.15 mL/min. The sample injection volume was 5 µL. ESI source parameters were as follows: spray voltage, 3200 V or −2800 V, in positive or negative modes, respectively; sheath gas, 35 arb (arbitrary units); aux gas, 10 arb; sweep gas, 0.5 arb; ion transfer tube temperature, 300 °C; vaporizer temperature, 35 °C. LC–MS data acquisition was operated under full scan polarity switching mode for all samples. The full scan was set as: orbitrap resolution, 120,000 at m/z 200; AGC target, 1e7; maximum injection time, 200 ms; scan range, 60–1000 m/z.

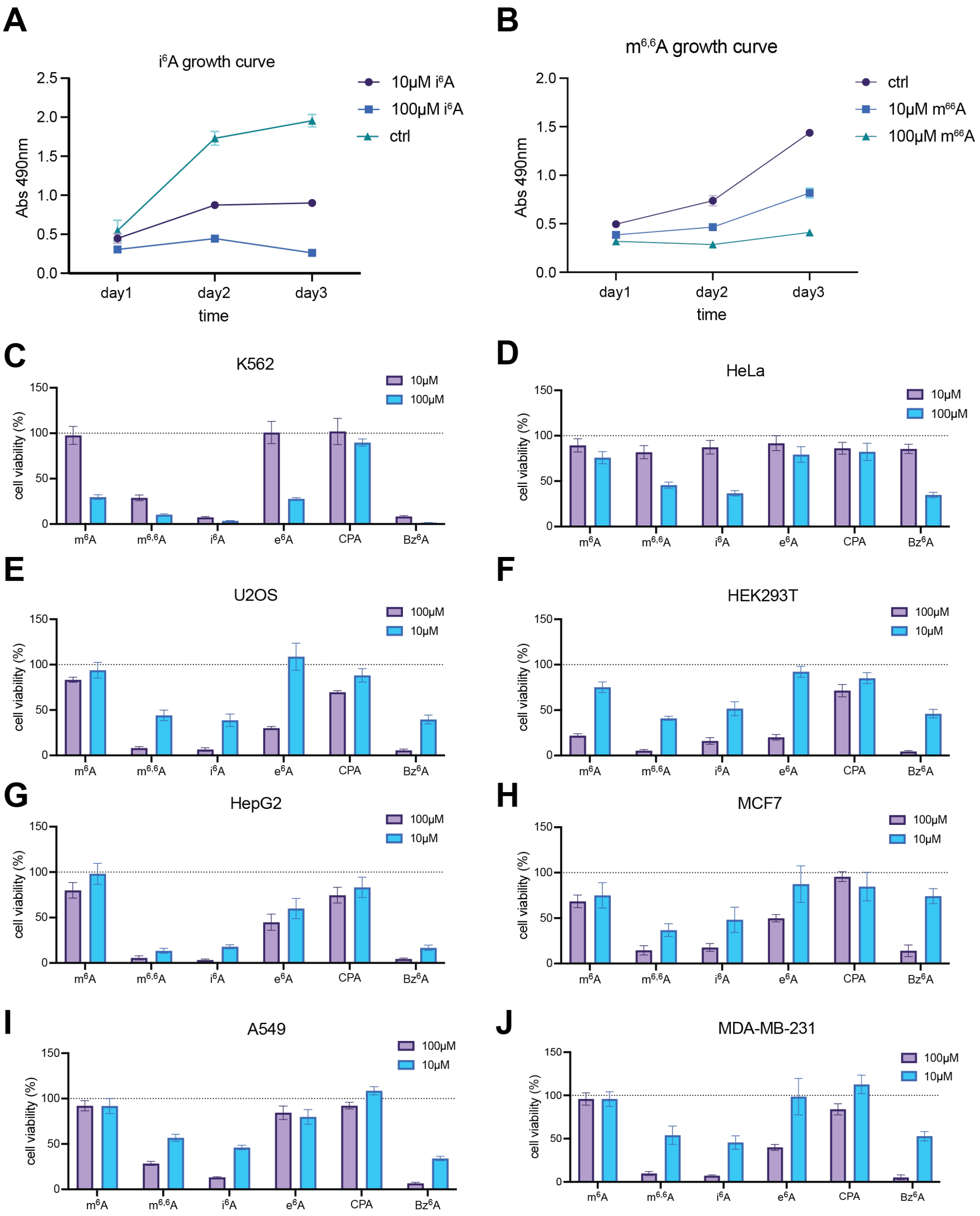


**Supplementary Figure 1.** MTS-based cell viability measurements of cytotoxicity caused by $N^6$-modified adenosine. **(A-B)** 3-day growth curves for A549 cells treated with 10 or 100μM $i^6A$ **(A)** or $m^{6,6}A$ **(B)**. Normalization was conducted by subtraction of no cell control and data are mean ± s. d. (*n=9* with three technical replicates for each of the three independent biological replicates). **(C-J)** Screening of $N^6$-modified adenosines cytotoxicity in a panel of human cell lines by treating 10 μM or 100 μM nucleosides for 72 h. Data corresponds to Figure 1E in the main text. Cell viability of K562 **(C)**, HeLa **(D)**, U2OS **(E)**, HEK293T **(F)**, HepG2 **(G)**, MCF7 **(H)**, A549 **(I)** and MDA-MB-

231 **(J)** was normalized by untreated cells and data are mean ± s. d. (*n=9* with three technical replicates for each of the three independent biological replicates).

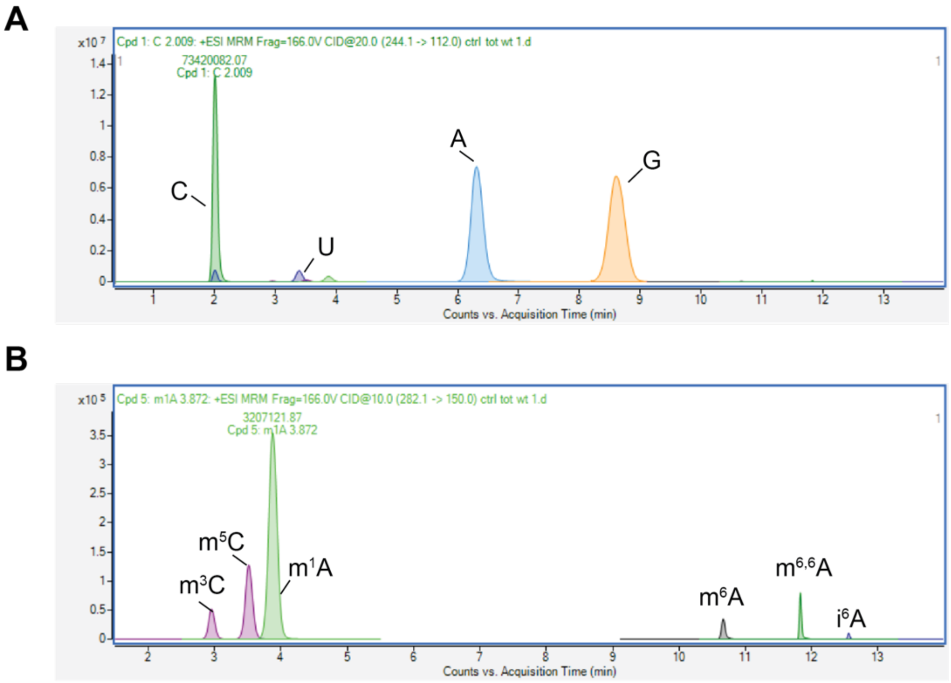


**Supplementary Figure 2.** Representative LC-QQQ-MS chromatograms for A, G, C, U **(A)** and $m^5C$, $m^1A$, $m^6A$, $m^{6,6}A$ and $i^6A$ **(B)** from total RNA of WT HEK293T. Ion counts were measured in DMRM mode.

Cytidine (C)
500 ng/mL to 5 ng/mL

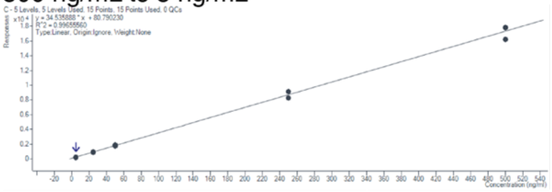

Uridine (U)
500 ng/mL to 5 ng/mL

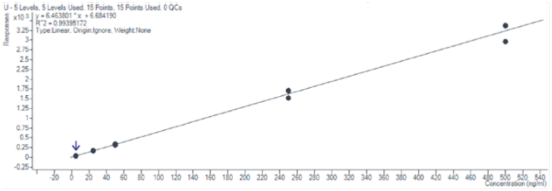

N1-methylAdenosine ($m^1A$)
50 ng/mL to 0.5 ng/mL

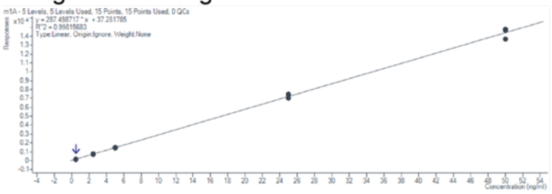

Adenosine(A)
500 ng/mL to 5 ng/mL

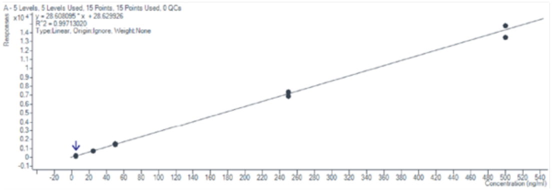

Guanosine(G)
500 ng/mL to 5 ng/mL

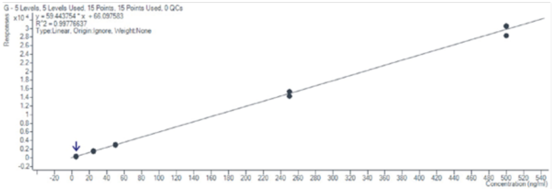

N6-methylAdenosine ($m^6A$)
50 ng/mL to 0.5 ng/mL

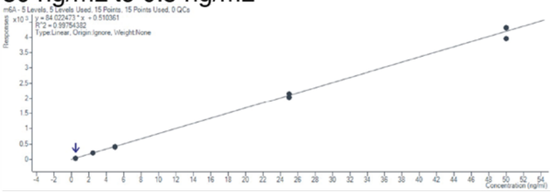

N6,N6-dimethylAdenosine ($m^{6,6}A$)
50 ng/mL to 0.5 ng/mL

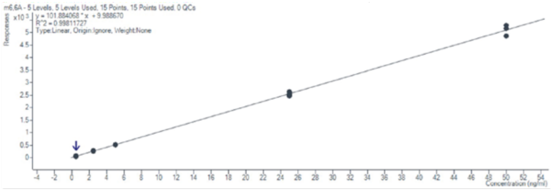

N6-cyclopentylAdenosine (CPA)
50 ng/mL to 0.5 ng/mL

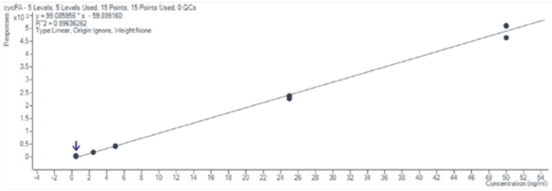

N6-isopentenylAdenosine ($i^6A$)
50 ng/mL to 0.5 ng/mL

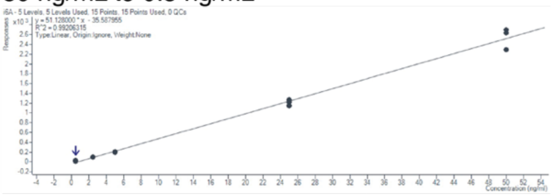

N6-benzylAdenosine ($Bz^6A$)
50 ng/mL to 0.5 ng/mL

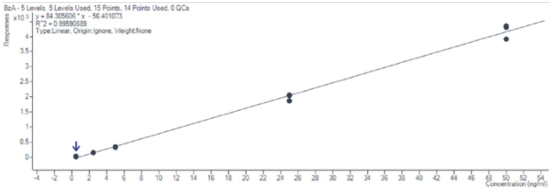

**Supplementary Figure 3.** Representative standard curves used for LC-QQQ-MS quantification of modified ribonucleosides. Two technical replicates were used to generate standard curves.

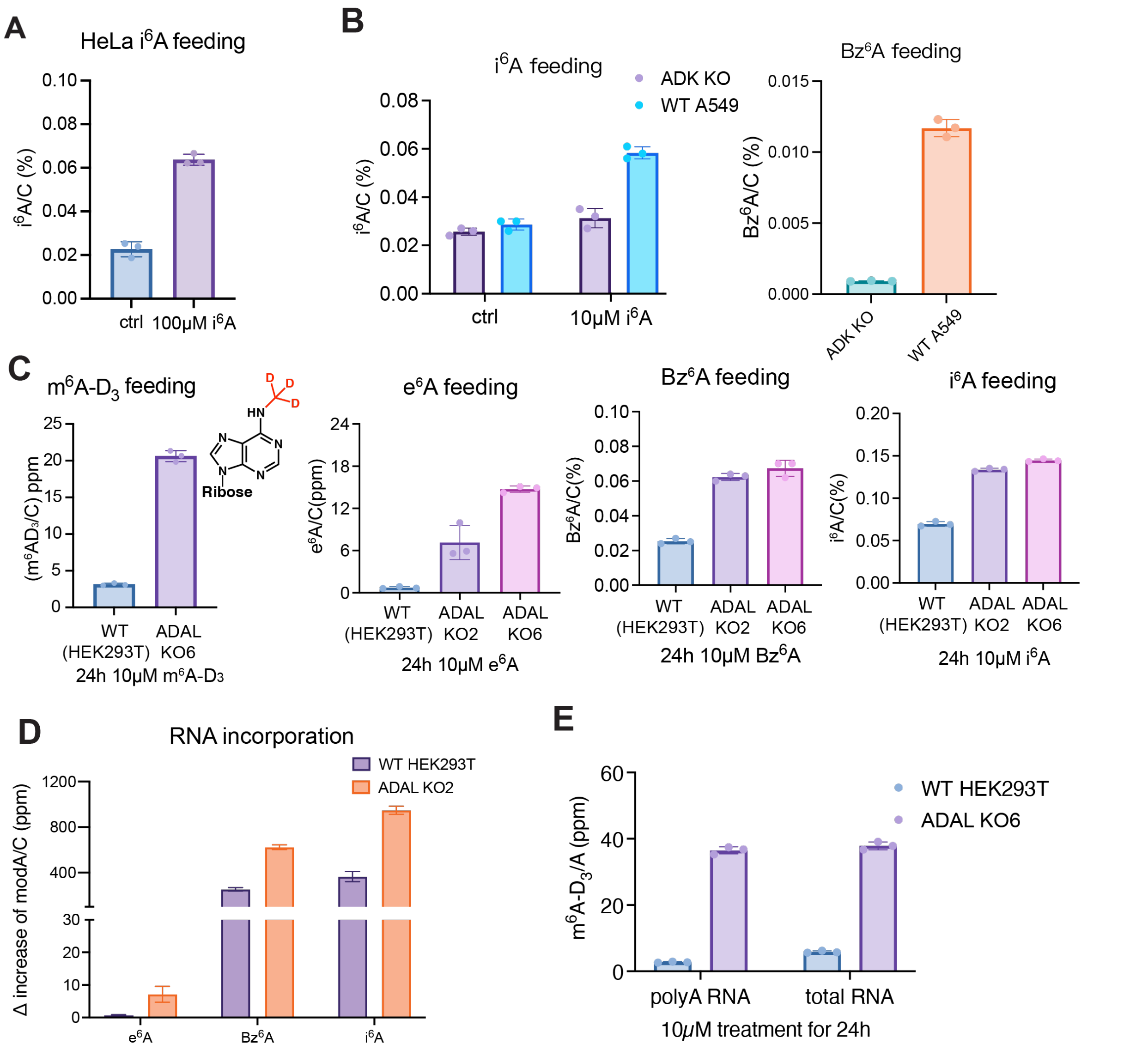


**Supplementary Figure 4.** RNA incorporation of $N^6$-modified adenosine analogs quantified by nucleoside LC-QQQ-MS. Three independent biological replicates were assayed. Data are mean ± s.d. **(A)** $i^6A$ incorporation rate in HeLa with 100μM $i^6A$ for 24h. **(B)** $i^6A$ or $Bz^6A$ incorporation rate in WT A549 or A549 ADK KO. Cells were fed with 10μM $i^6A$ or $Bz^6A$ for 24h. **(C)** $m^6A$-$D_3$, $e^6A$, $Bz^6A$ or $i^6A$ incorporation rate in WT HEK293T and ADAL KO2 or KO6. Cells were fed with 10μM nucleoside for 24h. **(D)** Δ increase of RNA modification levels in HEK293T WT and ADAL KO2 after 10μM nucleoside treatment for 24h. Δ increase was calculated from Supplementary Figure 4C and data are mean ± s.d. (*n*=3). **(E)** $m^6A$-$D_3$ incorporation rate in WT HEK293T and ADAL KO6. Cells were fed with 10μM nucleoside for 24h.

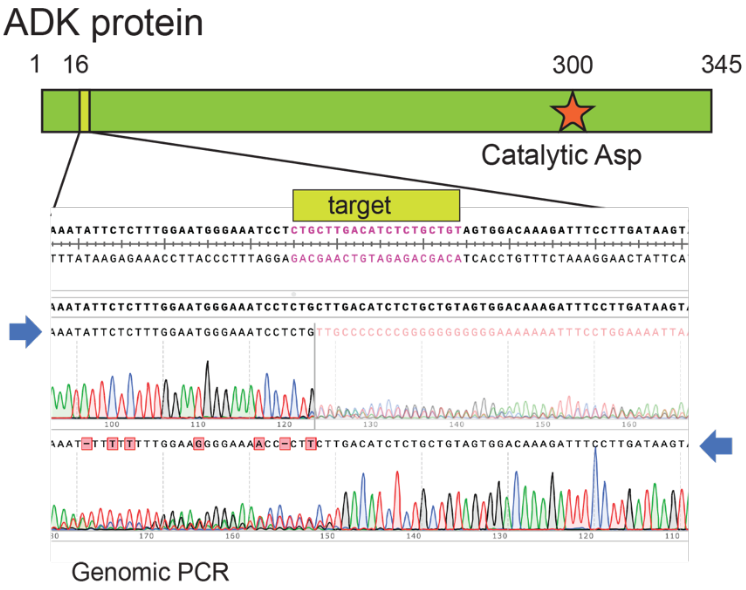


**Supplementary Figure 5.** Schematic of guide RNA targeting ADK gene and genomic PCR Sanger sequencing results for A549 ADK KO.

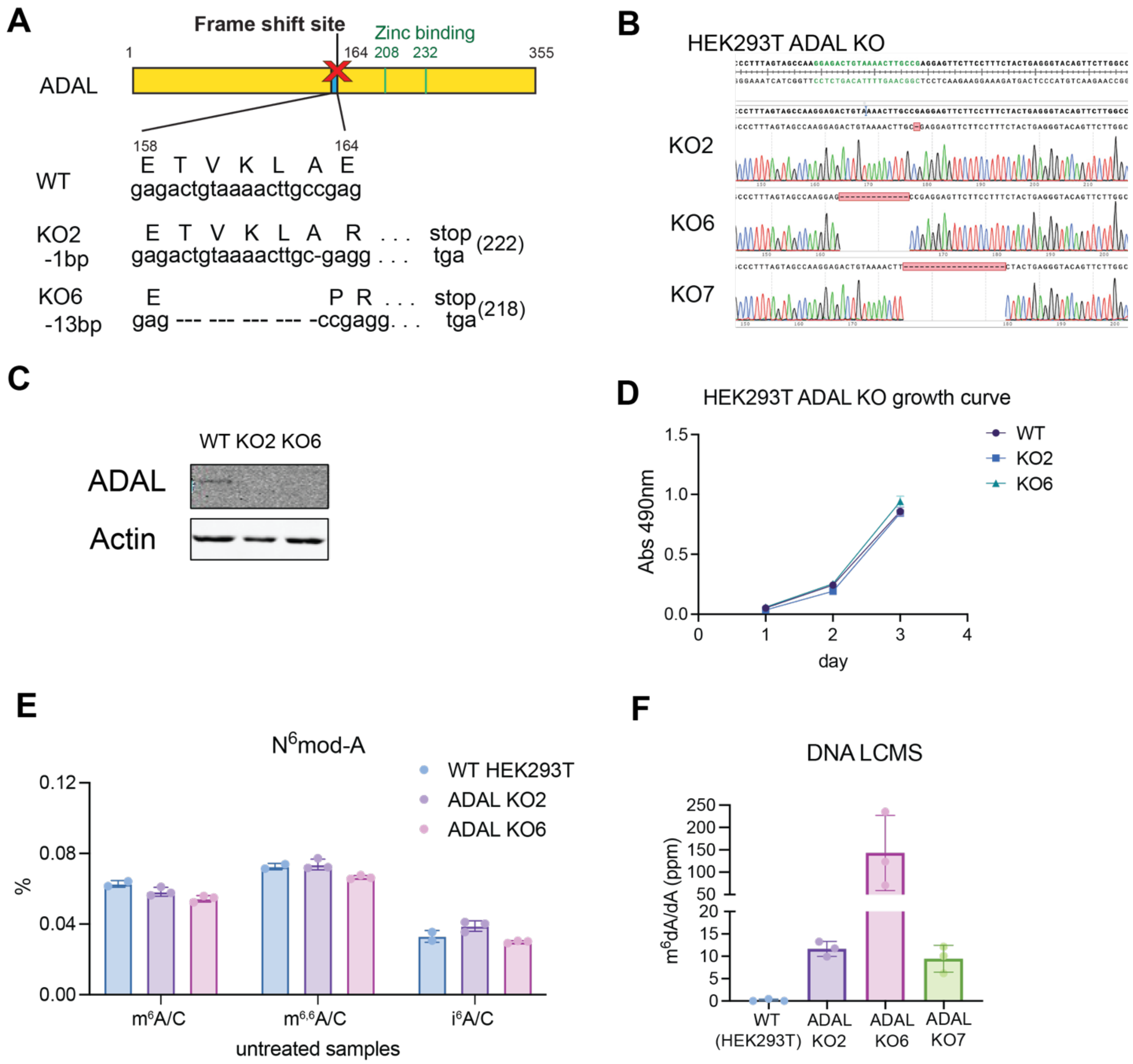


**Supplementary Figure 6.** Validation and characterization of ADAL KO in HEK293T. **(A)** Schematic cartoon of ADAL protein sequence and frame shift sequences in ADAL KO2 and ADAL KO6. **(B)** Genomic Sanger sequencing data for HEK293T ADAL KO2, KO6 and KO7 at guide RNA targeting location. **(C)** Western blot validation of ADAL KO in HEK293T cells. **(D)** 3-day growth curves of HEK293T WT, ADAL KO2 and KO6. Cell viability was measured by an MTS-based assay. Data are mean ± s. d. (*n*=9 with three technical replicates for each of the three independent biological replicates). **(E)** Endogenous abundance of $N^6$-modified adenosines in total RNA from HEK293T WT, ADAL KO2 and KO6 cells without exogenous nucleoside treatment. Modification levels were quantified by nucleoside LC-QQQ-MS. Data are mean ± s.d. (*n*=3). **(F)** Endogenous $N^6$-methyl-2'deoxyadenosine levels from genomic DNA in WT HEK293T, ADAL KO2, KO6 and KO7 cells without exogenous nucleoside treatment. Modification levels were quantified by nucleoside LC-QQQ-MS. Data are mean ± s.d. (*n*=3). Unpaired t-test was performed, and p-values are WT vs KO2: p=0.0003; WT vs KO6: p=0.042; WT vs KO7: p=0.006.

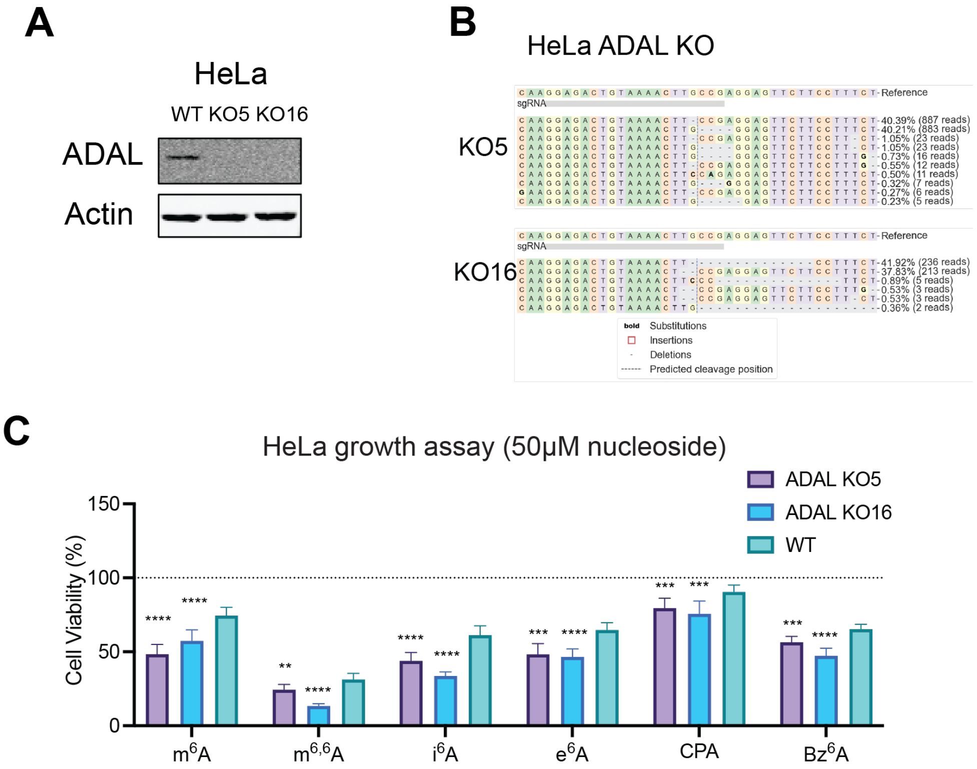


**Supplementary Figure 7.** Validation and characterization HeLa ADAL KO. **(A)** Western blot validation of ADAL KO in HeLa cells. **(B)** Sanger sequencing of sgRNA genomic targets sites in HeLa ADAL KO5 and KO16. **(C)** Quantification of cell growth in HeLa WT and ADAL KO5/KO16 cells using an MTS-based assay. Cells were treated with $N^6$-modified adenosine analogs at 50μM for 72h. Plot represents mean normalized cell viability ± s. d. (*n*=9 with three technical replicates for each of the three independent biological replicates). Multiple unpaired t-test were performed between WT and ADAL KOs. Adjusted p values: for $m^6A$: WT vs KO5: p<0.000001; WT vs KO16: p=0.00009; for $m^{6,6}A$: WT vs KO5 p=0.0019; WT vs KO16 p<0.000001; for $i^6A$: WT vs KO5: p=0.000059; WT vs KO16 p<0.000001; for $e^6A$: WT vs KO5 p=0.00014; WT vs KO16 p=0.000003; for CPA: WT vs KO5 p=0.00019; WT vs KO16 p=0.000331; for $Bz^6A$: WT vs KO5 p=0.00026; WT vs KO16 p<0.000001.

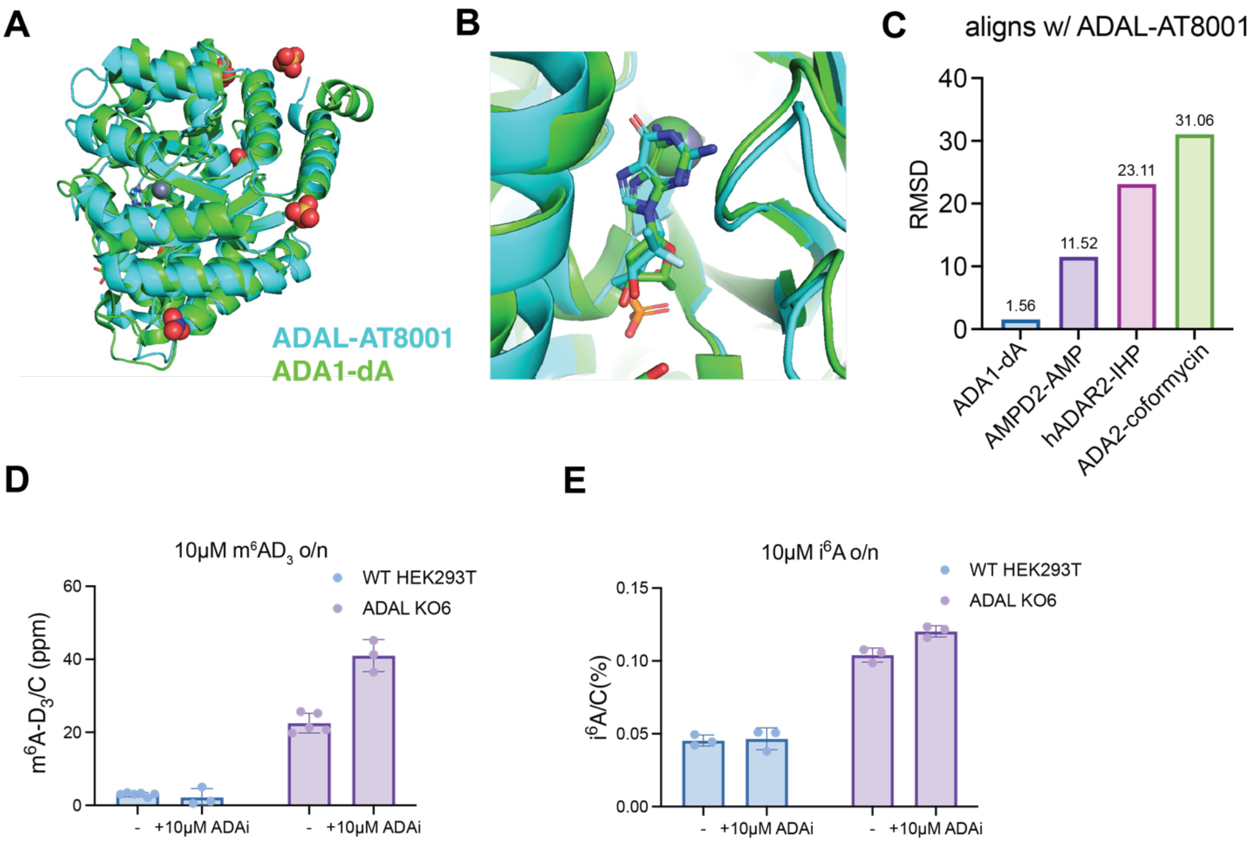


**Supplementary Figure 8.** ADA protein shares similar structure and function as ADAL protein. **(A)** Crystal structure alignment of ADAL (PDB: 8QCH) and ADA1 (PDB: 3IAR) by Pymol. **(B)** Close up view for binding pocket of structure alignment of ADAL (PDB: 8QCH) and ADA1 (PDB: 3IAR) by Pymol. **(C)** RMSD score of different alignment with ADAL (PDB: 8QCH). RMSD scores were calculated by Pymol. **(D-E)** RNA incorporation of $N^6$-adenosine analogs in HEK293T WT and ADAL KO6 with or without ADAi treatment (10μM Pentostatin). ADAi treated cells were pre-treated with 10μM pentostatin for 6 hours and then exposed to both nucleosides (10 μM) and pentostatin overnight. Cells without ADAi treatement were only incubated with 10μM overnight and harvested together with the treated samples. $m^6A$-$D_3$ **(D)** or $i^6A$ **(E)** levels in total RNA were quantified by nucleoside LC-QQQ-MS. Data are mean ± s.d. For $m^6A$-$D_3$ feeding, WT without ADAi: *n*=6; ADAL KO6 without ADAi: *n*=5; WT or ADAL KO6 with ADAi: *n*=3. Three independent biological replicates were assayed for $i^6A$ treated cells. Multiple unpaired t-test were performed between ADAi treated and untreated samples. Adjusted p-values for $m^6A$-$D_3$: p=0.00056; for $i^6A$: p=0.03.

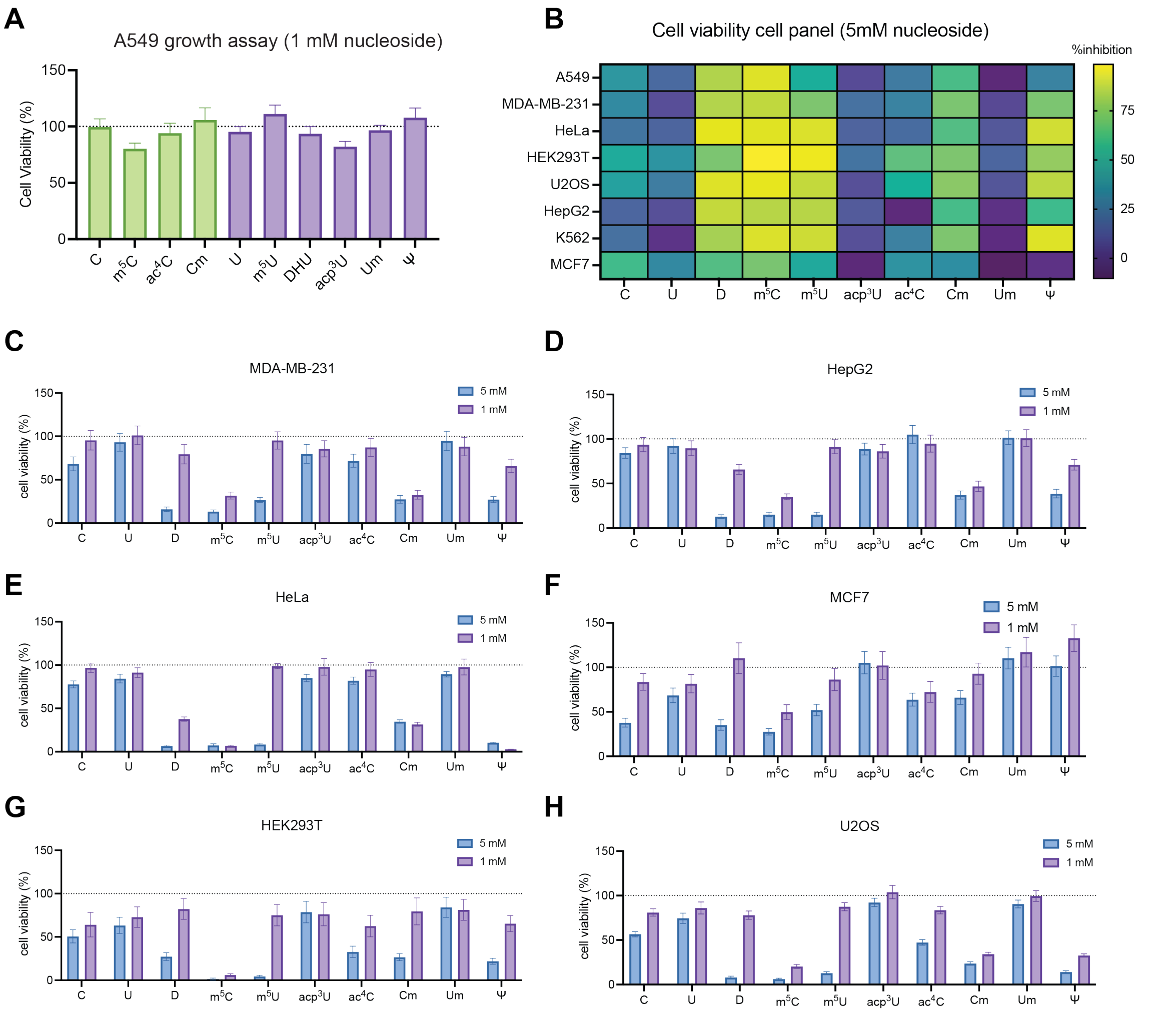


**Supplementary Figure 9.** Cytotoxicity of epigenetic ribopyrimidine analogs**. (A)** Screening in WT A549 cells at 1 mM (72h incubation time). Cell viability was measured by an MTS-based assay and plot represents mean normalized cell viability ± s. d. (*n*=9 with three technical replicates for each of the three independent biological replicates) **(B)** Heatmap shows percentage of cell growth inhibition across different cell lines with 5mM epigenetic pyrimidine analogs for 72h incubation time, corresponding to Figure 3B and Supplementary Figure 9C-H. Cell viability was measured by an MTS-based assay and mean normalized by untreated cells. (*n*=9 with three technical replicates for each of the three independent biological replicates) **(C-H)** Screening epigenetic ribopyrimidines cytotoxicity in a panel of human cell lines by treating 1 mM or 5 mM nucleosides for 72 h. Corresponding to the Figure 3C. Cell viability of MDA-MB-231 **(C)**, HepG2 **(D)**, HeLa **(E)**, MCF7 **(F)**, HEK293T **(G)** and U2OS **(H)** was normalized by untreated cells and data are mean ± s. d. (*n*=9 with three technical replicates for each of the three independent biological replicates).

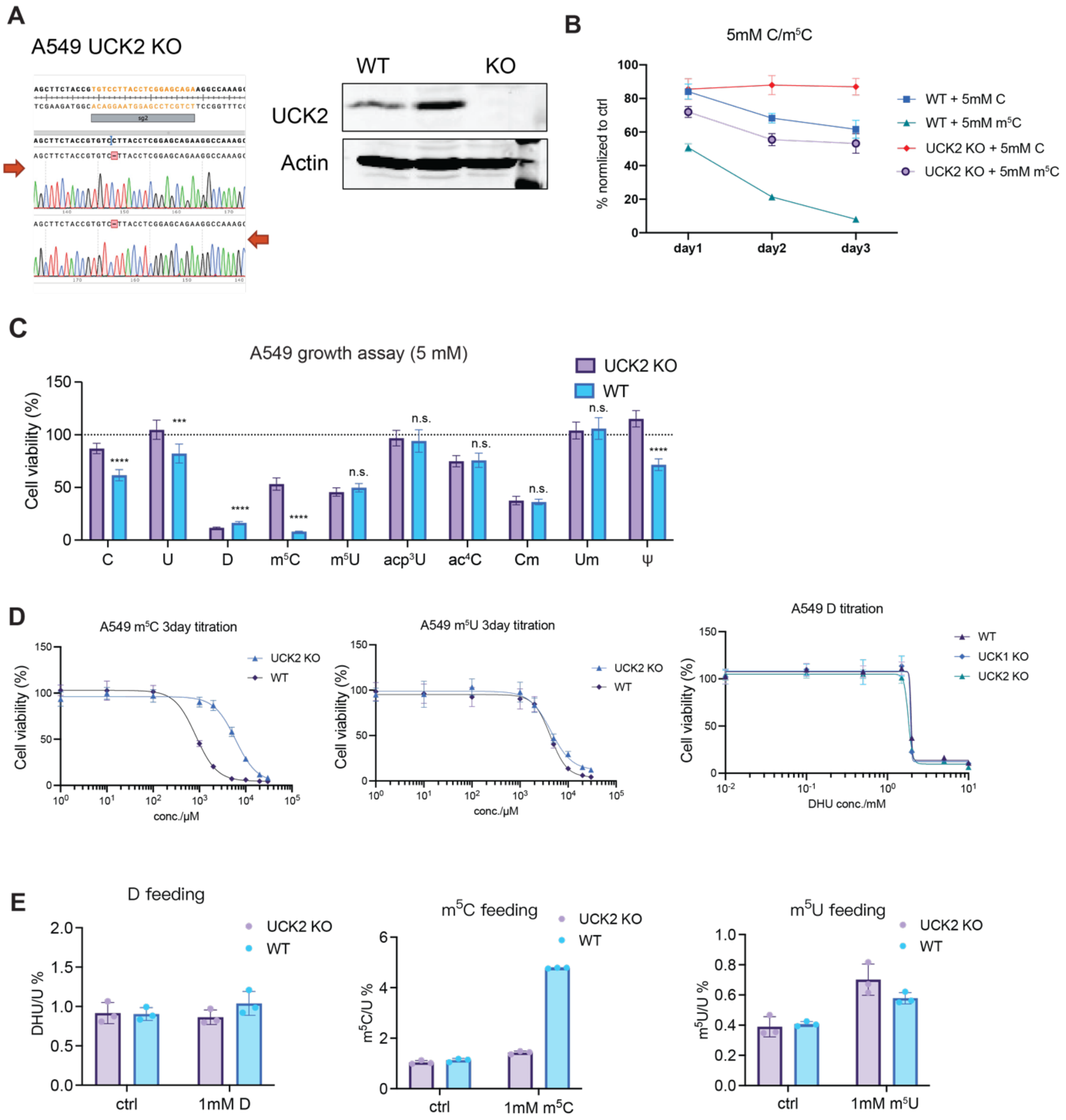


**Supplementary Figure 10.** UCK2 KO in A549 cells. **(A)** Validation of UCK2 KO in A549 by genomic PCR and Western blot. **(B)** 3-day cell viability measurements by an MTS-based assay from A549 WT and UCK2 KO with 5mM C or $m^5C$ treatment. Plot represents mean normalized cell viability ± s. d. (*n*=9 with three technical replicates for each of the three independent biological replicates). **(C)** Cytotoxicity of different epigenetic pyrimidine analogs (5 mM treatment for 72 hr) in A549 WT and UCK2 KO cells. Cell viability was measured by an MTS-based assay and plot represents mean normalized cell viability ± s. d. (*n*=9 with three technical replicates for each of the three independent biological replicates) **(D)** Titration curves for pyrimidine analogs in A549. $IC_{50}$ values of $m^5C$ titrations: WT: 835.8 μM; UCK2 KO: 6020 μM. $IC_{50}$ values of $m^5U$ titrations: WT: 4281μM; UCK2 KO: 4365 μM. Cell viability was measured by an MTS-based assay and plot represents mean normalized cell viability ± s. d. Curves were fitted based on a 4-parameter dose-response equation using GraphPad Prism. (*n*=9 with three technical replicates for each of the three independent biological replicates) **(E)** RNA modification levels in total RNA quantified by nucleoside LC-QQQ-MS after feeding with 1mM D, $m^5C$ or $m^5U$ in A549 WT and UCK2 KO cells for 24h. Data are mean ± s.d. (*n*=3).

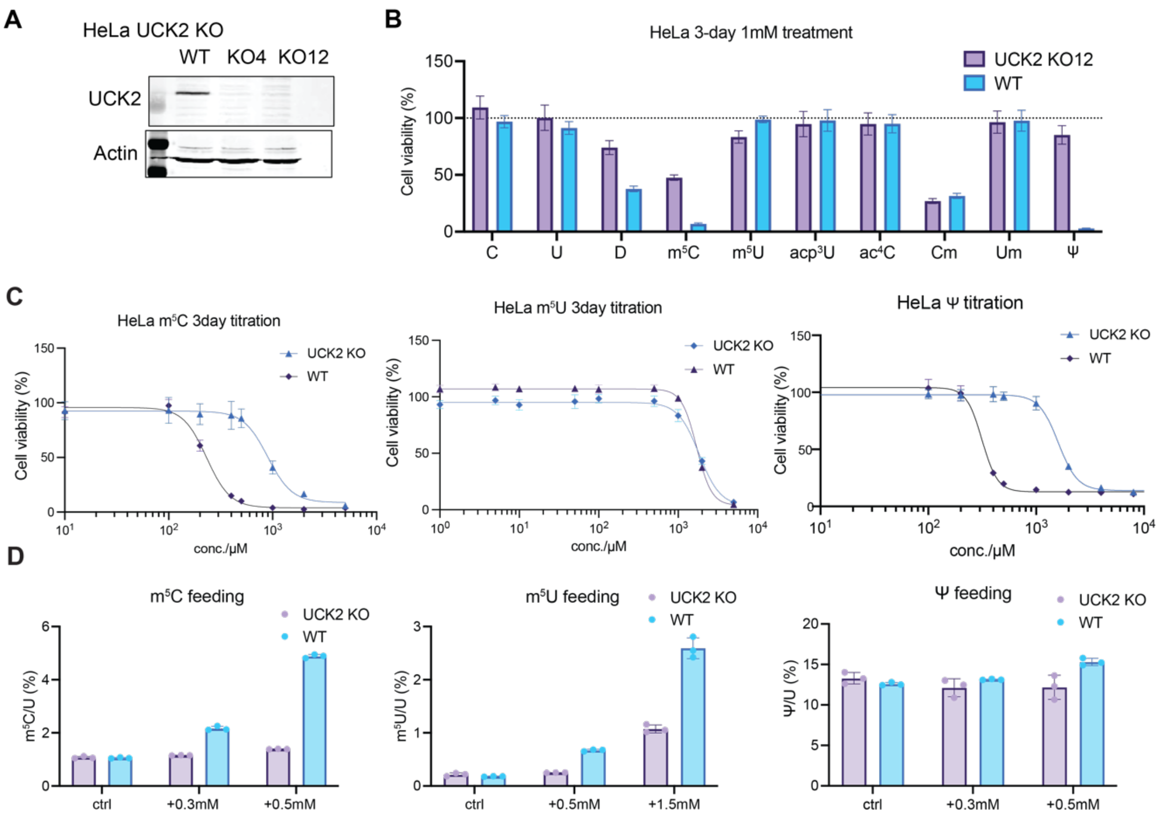


**Supplementary Figure 11.** UCK2 KO in HeLa cells. **(A)** Validation of UCK2 KO in HeLa by Western blot. **(B)** Cytotoxicity of epigenetic ribopyrimidine analogs (1 mM for 72 hr) in HeLa WT and UCK2 KO cells. Cell viability was measured by an MTS-based assay and plot represents mean normalized cell viability ± s. d. (*n*=9 with three technical replicates for each of the three independent biological replicates) **(C)** Titration curves for pyrimidine analogs in HeLa. $IC_{50}$ values of $m^5C$ titrations: WT: 231.6 μM; UCK2 KO: 897.6 μM. $IC_{50}$ values of $m^5U$ titrations: WT: 1708 μM; UCK2 KO: 1839 μM. $IC_{50}$ values of Ψ titrations: WT: 314.6μM; UCK2 KO: 1605μM. Cell viability was measured by an MTS-based assay and plot represents mean normalized cell viability ± s. d. Curves were fitted based on a 4-parameter dose-response equation using GraphPad Prism. (*n*=9 with three technical replicates for each of the three independent biological replicates) **(D)** RNA modification levels quantified by nucleoside LC-QQQ-MS upon feeding with 0.3 or 0.5 mM $m^5C$ or Ψ, and 0.5 or 1.5 mM $m^5U$ in HeLa WT and UCK2 KO cells for 24h. Data are mean ± s.d. (*n*=3).

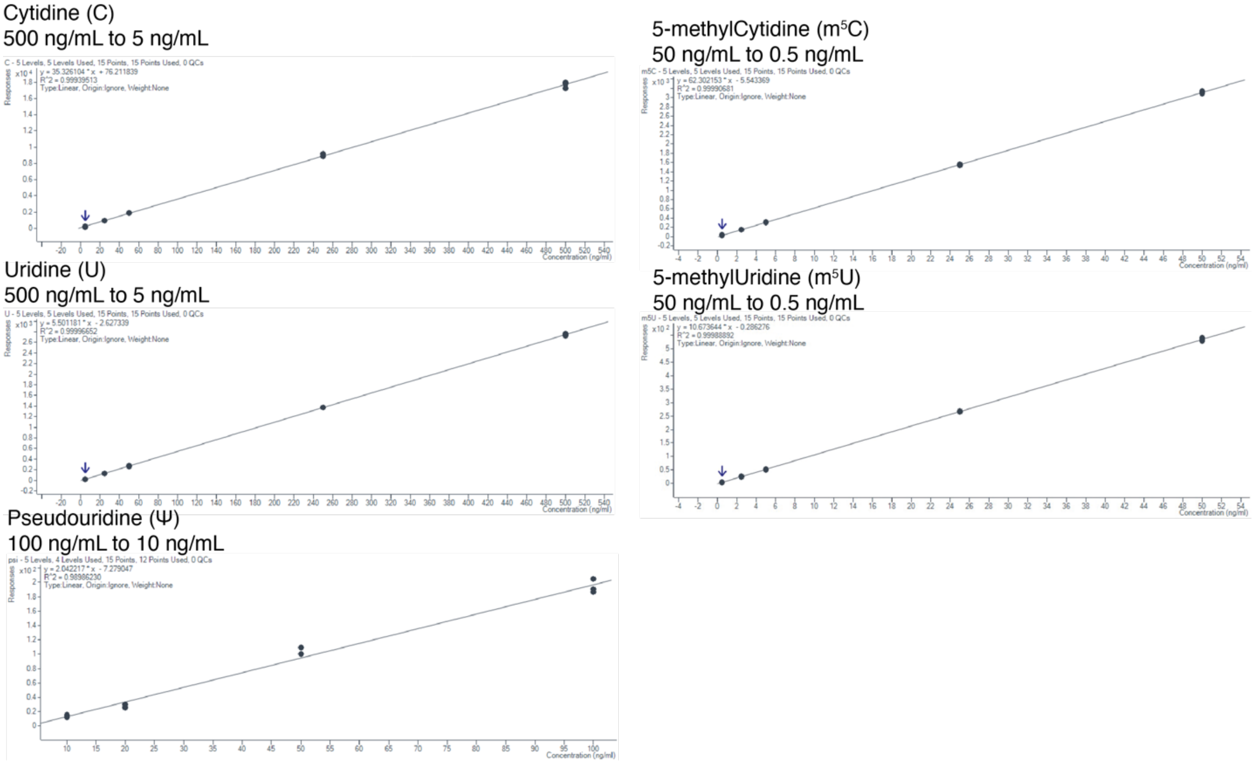


**Supplementary Figure 12.** Representative standard curves used for LC-QQQ-MS quantification of pyrimidine analogs. Two technical replicates were used to generate standard curves.

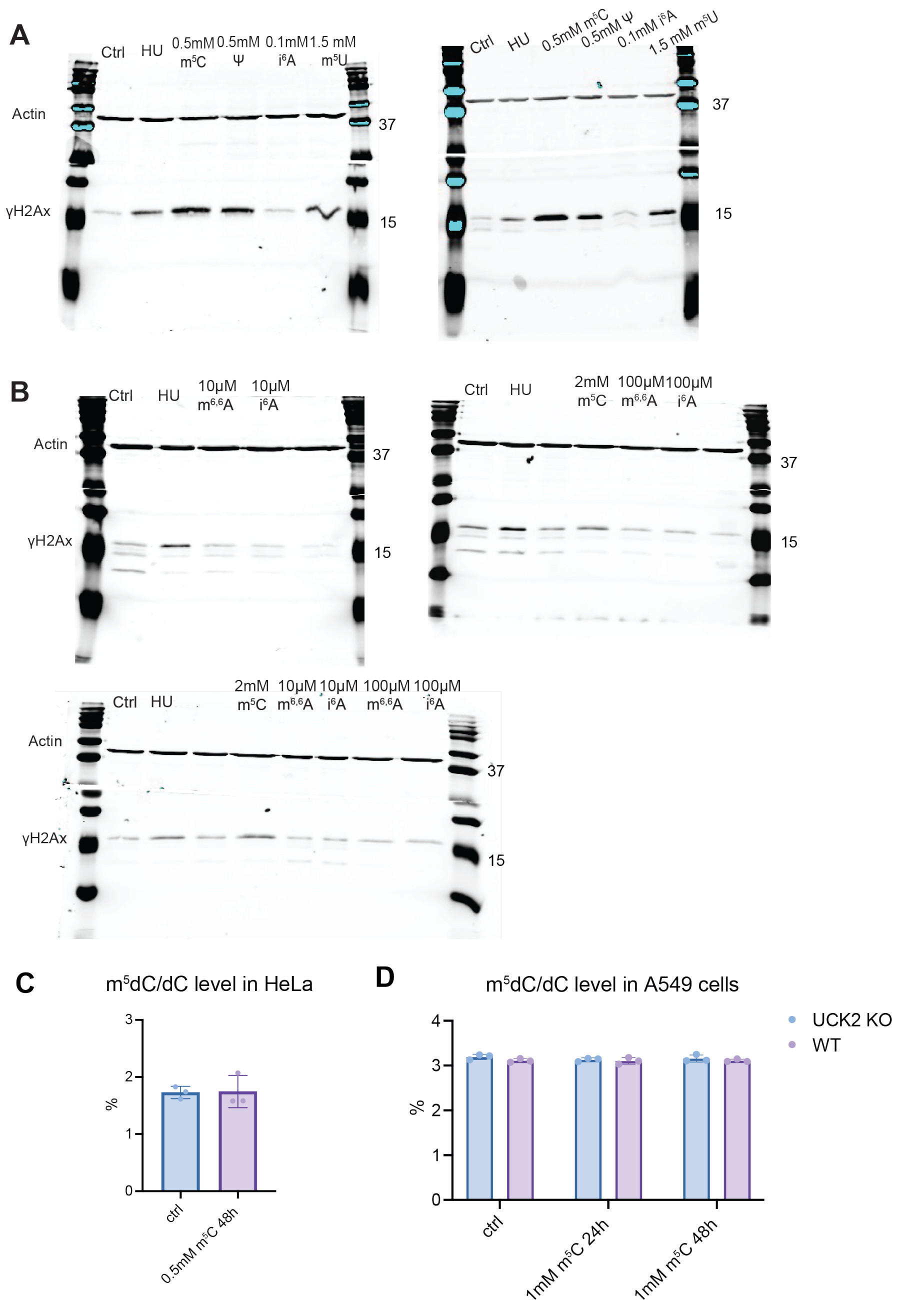


**Supplementary Figure 13.** DNA damage and DNA incorporation analysis upon nucleoside treatment. **(A)** γH2Ax Western blot of lysate from HeLa cells treated with 10 mM Hydroxyurea (HU, 6h), 0.5 mM $m^5C$ (48h), 0.5 mM Ψ (48h), 0.1 mM $i^6A$ (48h) or 1.5 mM $m^5U$ (48h). Actin served as loading control. **(B)** γH2Ax Western blot of lysate from A549 cells treated with 10 mM Hydroxyurea (HU, 6h), 2 mM $m^5C$ (48h), 10 μM or 100 μM $m^{6,6}A$ or $i^6A$ (48h). Actin served as loading control. **(C)** $m^5dC$ quantified by nucleoside LC-QQQ-MS in HeLa cells fed with 0.5 mM $m^5C$ for 48 h. Data are mean ± s. d. (*n*=3) **(D)** $m^5dC$ quantified by nucleoside LC-QQQ-MS in A549 cells fed with 1 mM $m^5C$ for 24 h or 48 h. Data are mean ± s. d. (*n*=3)

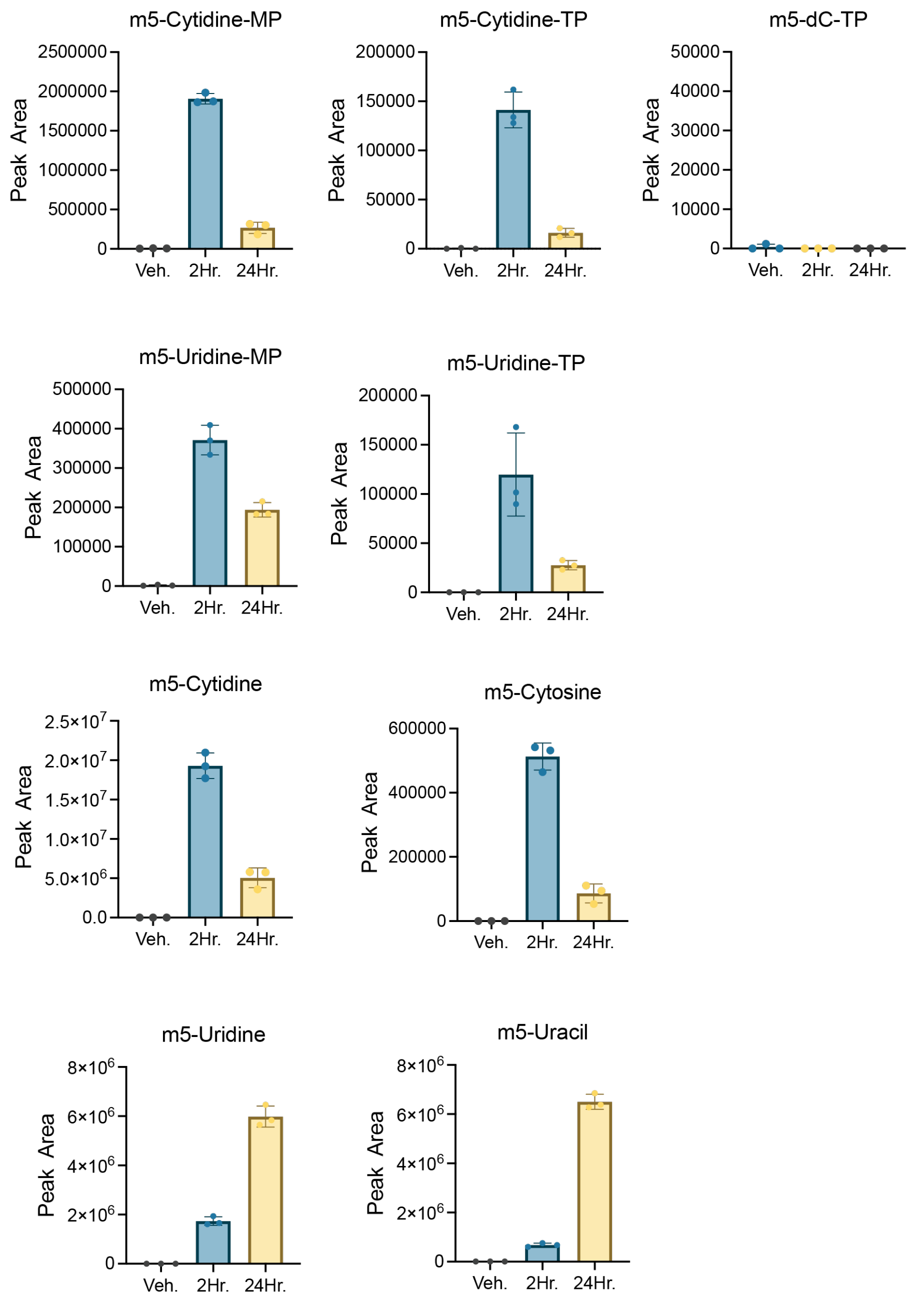


**Supplementary Figure 14.** LC-MS analysis of nucleotide pools in WT HeLa treated with 500 μM $m^5C$ for 2 h or 24 h. Peak area of 5-methylcytidine monophosphate ($m^5CMP$), 5-methylcytidine triphosphate ($m^5CTP$), 5-methyl-2'-deoxycytidine triphosphate ($m^5dCTP$), 5-methyluridine monophosphate ($m^5UMP$), 5-methyluridine triphosphate ($m^5UTP$), $m^5$-cytidine ($m^5C$), $m^5$-cytosine, $m^5$-uridine ($m^5U$) and $m^5$-uracil (thymine) were plotted. Data are mean ± s.d. (*n*=3).

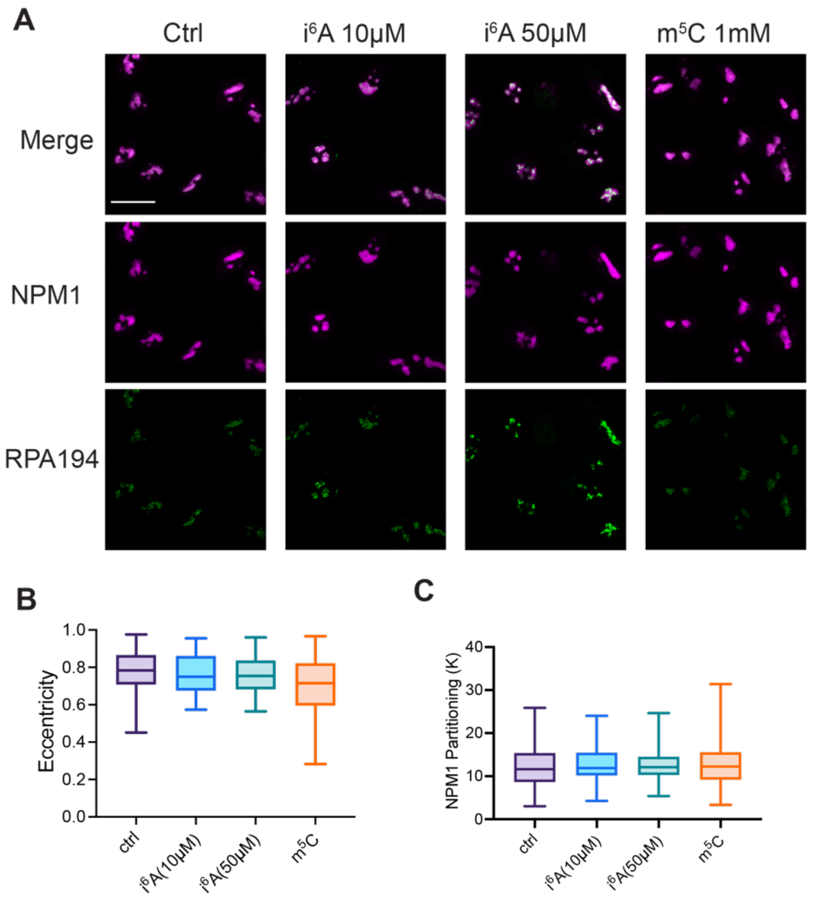


**Supplementary Figure 15.** Nucleolar stress induced by $i^6A$ in WT A549 cells. **(A)** Confocal images of WT A549 treated with 10 μM/50 μM $i^6A$ or 1 mM $m^5C$ for 24 h with NPM1 and RPA-194 staining. **(B)** Quantification of nucleolar eccentricity. **(C)** Quantification of NPM1 partitioning. Boxes in the plot represent $25^{th}$ to $75^{th}$ percentiles and whiskers show min to max. Representative images from 2 biological replicates. Between 40-120 cells were analyzed per condition.

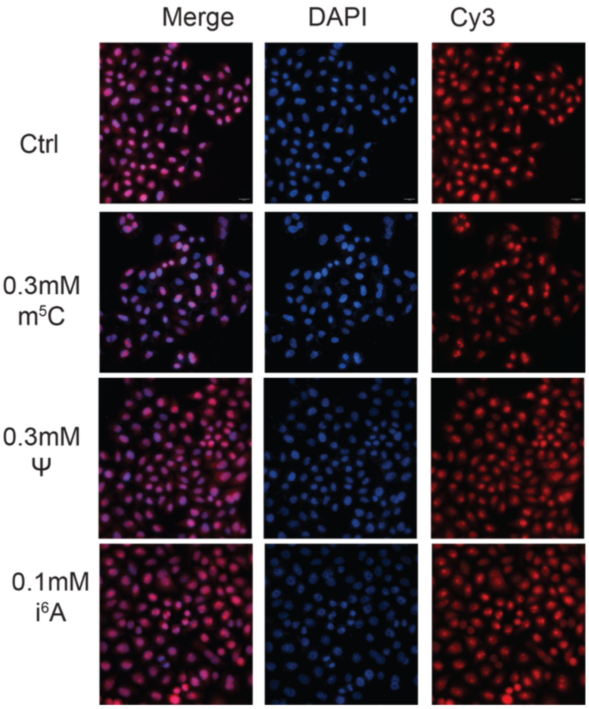


**Supplementary Figure 16.** Global translation levels upon nucleoside feeding by OP-Puro assay. Representative microscope images of WT HeLa treated with 100 µM $i^6A$, 500 µM $m^5C$ or 500 µM Ψ for 24 h. Images correspond to Figure 4D.
